# Statins and genetic inhibition of the mevalonate pathway activate an ATF3-STMN2 regenerative program

**DOI:** 10.64898/2026.02.23.707492

**Authors:** Matthew Nolan, Sandeep Aryal, I. Sandra Ndayambaje, Maize Cao, Peyton Lee, Moriah Hovde, Shuqi Yun, Josette Wlaschin, Aaron Held, Hortense Beaussant, Benjamin Wymann, Chao Zong-Lee, Su Min Lim, Xin Jiang, Nandini Ramesh, Ana Rita Agra Almeida Quadros, Ayub Boulos, Nicolas Zinter, Shireen Salem, Laïla El-Tayar, Melinda Beccari, Maximiliano Presa, Charles Jourdan Ferreras Reyes, Yin Yin Ruan, Grant Griesman, Corey Aguilar, James Hawrot, Hayden Wheeler, Zevik Melamed, Benjamin P. Kleinstiver, Mark Albers, Don W. Cleveland, Rudolph E. Tanzi, Cathleen M. Lutz, Robert D. Hubbard, Dione Kobayashi, Michael Ward, Christiano R.R Alves, Brian Wainger, Claire Le Pichon, Clotilde Lagier-Tourenne

**Author notes:** **Corresponding author:** Clotilde Lagier-Tourenne.

## Abstract

Loss of neuronal regenerative capacity is a common feature of neurodegenerative disease and axonal injury, yet the transcriptional programs governing this state remain poorly defined. Stathmin-2 (STMN2), a tubulin-binding protein essential for axon maintenance and repair, is profoundly depleted following loss of nuclear TDP-43 in neurodegenerative disease. Here, we identify statins as potent inducers of STMN2 expression. Pharmacological and genetic suppression of the mevalonate pathway, and subsequent prevention of protein geranylgeranylation, restored STMN2 levels in TDP-43 deficient cells and promoted neurite growth. STMN2 induction was abrogated when using a statin analogue unable to interact with HMG-CoA reductase, and through co-administration of mevalonate or geranylgeranyl diphosphate substrates. RNA-seq revealed that statins induce a coordinated pro-regenerative transcriptional response, including activation of the AP-1 transcription factor complex gene, *ATF3*. Loss of ATF3 attenuated STMN2 induction *in vitro*, and diminished injury-induced *Stmn2* upregulation in spinal motor neurons *in vivo*. These results demonstrate statins as modulators of ATF3 and STMN2 expression and highlight their therapeutic potential in neurodegenerative disease.

## Introduction

Abnormal cytoplasmic accumulation and nuclear clearance of TAR DNA-binding protein 43 kDa (TDP-43) in neurons and glia is the primary pathological feature in most cases of amyotrophic lateral sclerosis (ALS-TDP) and frontotemporal dementia (FTD-TDP)^1–3^, all cases of limbic-predominant age-related TDP-43 encephalopathy (LATE)^4^, and as a co-pathology in up to 50% of Alzheimer’s disease (AD) cases^5^. Loss of nuclear TDP-43 function is an early disease mechanism in TDP-43 proteinopathies and can be observed even without apparent aggregation pathology^6–8^. TDP-43 is a nuclear RNA-binding protein critical for the splicing, transport and translation of thousands of mRNAs^9–13^, however the transcript most sensitive to TDP-43 loss-of-function encodes the tubulin binding protein, stathmin-2 (STMN2)^14,15^. Loss of TDP-43 induces the expression of a cryptic exon within the first *STMN2* intron, eliciting premature stop codon formation, polyadenylation of the transcript, and an associated reduction in full-length protein expression^14,15^. *STMN2* misprocessing is now recognized as a pathological hallmark of postmortem tissue from patients with TDP-43 proteinopathy, including sporadic and familial forms of ALS^14,15^, FTD^16,17^, and patients with LATE or AD^18–20^. STMN2 is proposed to stimulate microtubule dynamics by promoting their disassembly^21^ through the sequestration of tubulin dimers in the growth cone^22,23^. Additionally, increased expression of STMN2 enhances neurite outgrowth^24,25^, and overexpression of STMN2 alone is sufficient to rescue axonal regenerative potential following injury in induced pluripotent stem-cell (iPSC)-derived motor neurons with TDP-43 depletion^14^. We previously demonstrated that antisense oligonucleotides (ASOs) sterically-binding the cryptic exon were efficient in restoring full-length STMN2 expression and promoting axonal recovery in TDP-43 depleted motor neurons^26^. Notably, depletion of *Stmn2* in mice reduces axonal diameter, elicits denervation of the neuromuscular junctions, and results in dose-dependent motor deficits - mimicking both pathological and clinical aspects of human ALS^27–29^. Together, these findings highlight STMN2 as a pertinent therapeutic target and suggest that modulation of its expression may be beneficial across a spectrum of neurodegenerative diseases.

Statins are inhibitors of β-Hydroxy β-methylglutaryl-CoA reductase (HMG-CoA reductase), and are commonly prescribed to treat hypercholesterolemia by preventing the conversion of HMG-CoA to mevalonate and thus reducing the production of low-density lipoprotein (LDL) cholesterol^30,31^. Beside their hypocholesterolemic action, statins exhibit pleiotropic effects that have been associated with beneficial outcomes in neurological disease^32^. The mevalonate pathway is also essential for the prenylation of proteins through the covalent binding of farnesyl pyrophosphate (FPP) or geranylgeranyl pyrophosphate (GGPP) moieties, catalyzed by the enzymes geranylgeranyl transferase (GGTase) I/II and farnesyl transferase (FTase). As upstream inhibitors of the mevalonate pathway, statins prevent the addition of these isoprenoid lipid modifications by decreasing the availability of FPP and GGPP substrates^33–35^. Prenylation is required for the correct subcellular localization, membrane interaction, and functioning of Rho-GTPases^36,37^. Rho-GTPase signaling mediates a variety of cellular processes including division, cytoskeletal organization, and intracellular trafficking, as well as transcription regulation via an upregulation of c-Jun and the AP-1 transcription factor complex^38,39^. In neurons, RhoA constrains axon growth through the condensation of actin and subsequent prevention of microtubule protrusion into the growth cone^40,41^, and previous studies have reported a pro-regenerative effect of statins both *in vitro*^42–46^ and *in vivo*^42,47^.

Here, we identify that chemical and genetic inhibition of the mevalonate pathway positively regulates STMN2 expression and neurite outgrowth. We demonstrate that statin-mediated induction of STMN2 results from the inhibition of geranylgeranylation and transcriptional upregulation of the AP-1 transcription factor complex, including ATF3. These results offer new insights into the regulation of STMN2 and provides evidence for therapeutic targeting of the mevalonate pathway in patients with TDP-43 proteinopathy.

### High-throughput screening identifies statins as potent activators of STMN2 expression

To uncover chemical modulators that increase STMN2 levels, we first generated luciferase reporter lines of STMN2 expression by tagging the N-terminus of both endogenous *STMN2* alleles with NanoLuciferase (*STMN2*-NLuc) in wild-type (WT) and TDP-43 mutant (N352S) SH-SY5Y cells **(Extended data Fig. 1a.)**. Immunoblot **(Fig. 1a)**, PCR **(Extended data Fig. 1b.)**, RT-PCR **(Extended data Fig. 1c.)**, and sequencing confirmed homozygosity and correct insertion of the tag. As expected, addition of the NanoLuciferase tag did not alter the sensitivity of *STMN2* pre-mRNA processing to TDP-43 loss **(Extended data Fig. 1d.)**. These engineered lines were used to develop a robust luciferase assay yielding NLuc signal proportional to cell densities **(Extended data Fig. 1e.)** and hygromycin-induced cell death **(Extended data Fig. 1f.)**. TDP-43 mutant SH-SY5Y cells display a significant reduction in *STMN2* mRNA and protein levels compared to WT^14^, a difference that was maintained after editing as determined by immunoblot and luciferase assays **(Fig. 1a, 1b.)**. Luciferase expression in the TDP-43 mutant line was subsequently used to screen 11,875 unique compounds spanning nine small-molecule libraries of both FDA-approved and experimental drugs **(Extended data Fig. 1g; Supplementary Table 1)**. TDP-43 mutant cells were chosen as the screening model to identify compounds that may return STMN2 expression to WT baseline despite TDP-43 dysfunction. TDP-43 mutant *STMN2*-NLuc cells were treated for 24 hours in duplicate at final concentrations ranging from 0.16-20μM depending on the format of the library. Median Z-prime score across the screen was 0.51 and Z-scores of technical replicates were highly correlated **(Extended data Fig. 1h.)**. Sixty-four compounds (0.53% of screened molecules) exhibited an average Z-score >3.0 in both technical replicates **(Extended data Fig. 1g-i)**, among which 52 compounds were selected for dose-response validation **(Table S2)**. Of these, six of the ten most effective molecules were statins, with cerivastatin being the most potent and yielding an EC50 of 0.08μM **(Fig. 1c; Supplementary Table 2)**.

**Figure 1.**
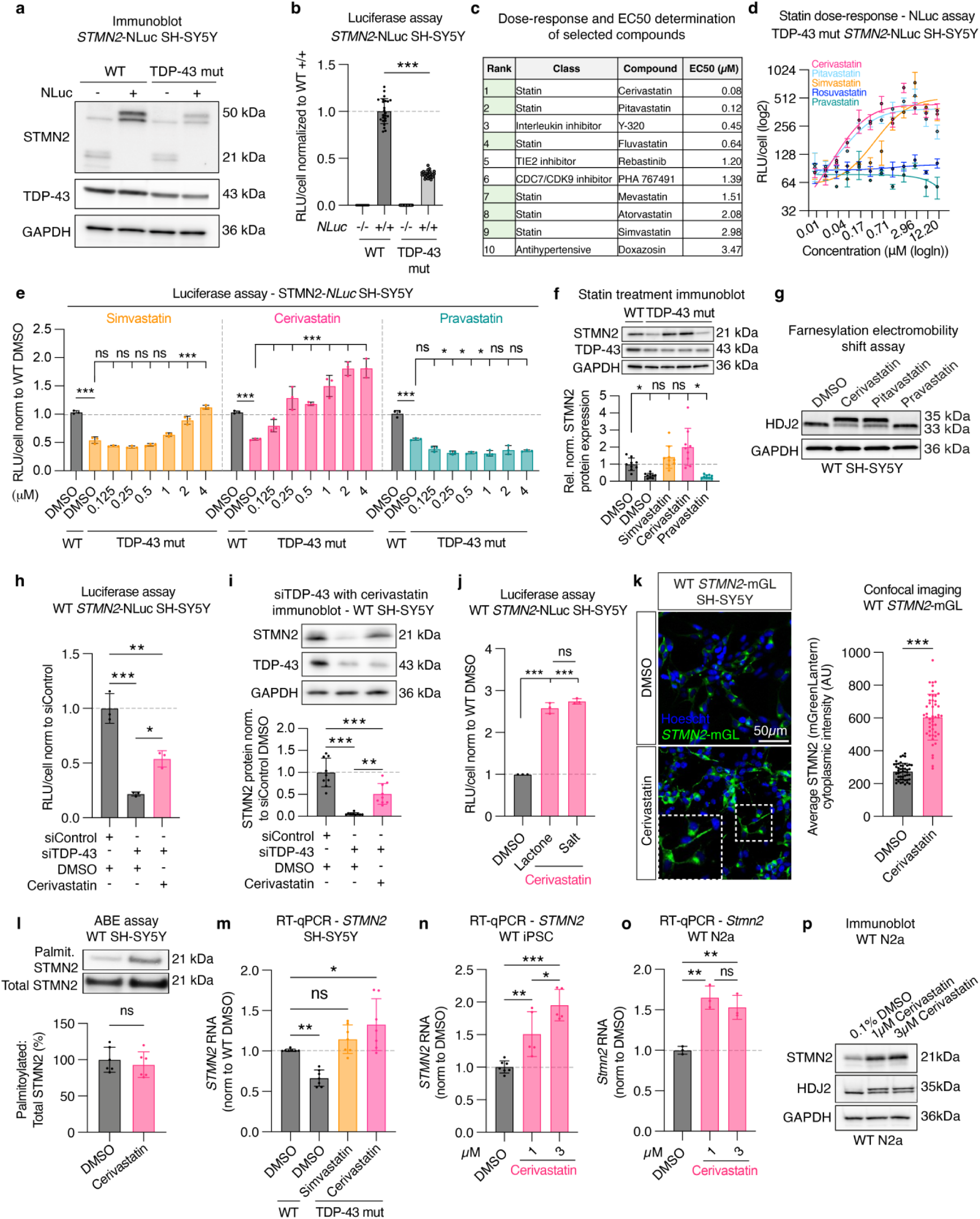
Pharmacological screen identifies statins as enhancers of STMN2 expression independently of TDP-43 function. **(A)** Immunoblot with antibodies against STMN2 and TDP-43 in WT and TDP-43 mutant (N352S) SH-SY5Y cells edited to express a Nanoluciferase tag (NLuc) fused to the STMN2 C-terminus on both endogenous alleles. GAPDH used as loading control. **(B)** Luciferase assay in edited lines measuring STMN2 protein level in TDP-43 mutant compared to WT cells. Points represent technical replicates from three independent experiments. One-way ANOVA with Tukey’s multiple comparisons correction. **(C)** Top 10 compounds identified through screening of 11,895 small molecules and validated by dose-response analysis. EC50 of each compound as listed. **(D)** Statin dose-response in TDP-43 mut *STMN2*-NLuc SH-SY5Y demonstrates that lipophilic (cerivastatin, simvastatin, pitavastatin), but not hydrophilic (rosuvastatin, pravastatin) statins promote STMN2 expression. Points represent mean of technical replicates in three independent experiments. Error bars = SEM. **(E)** Nanoluciferase assay in WT and TDP-43 mutant SH-SY5Y demonstrates that simvastatin and cerivastatin return STMN2 expression to WT baseline at ∼2μM and 0.25μM respectively. Points represent mean of technical replicates from three independent experiments. Two-way ANOVA with Tukey’s multiple comparisons correction. **(F)** Immunoblot and quantification of STMN2 protein in WT and TDP-43 mutant SH-SY5Y treated with DMSO, 2μM simvastatin, 1μM cerivastatin, or 3μM pravastatin. Points represent technical replicates from three independent experiments. GAPDH used as loading control. Kruskal-Wallis test, asterisks indicate comparisons to DMSO control. **(G)** Immunoblot for the heat-shock protein HDJ2 demonstrates that the lipophobic statins cerivastatin and pravastatin, but not pravastatin, inhibit farnesylation (demonstrated by a migration shift) in SH-SY5Y cells. GAPDH used as loading control. **(H)** Luciferase assay shows significant increase in STMN2 expression upon 1μM cerivastatin treatment of WT-SH-SY5Y *STMN2*-NLuc with siRNA-mediated TDP-43 suppression. Points represent mean of technical replicates from three independent experiments. One-way ANOVA with Tukey’s multiple comparisons. **(I)** Immunoblot with STMN2 and TDP-43 antibodies under same conditions as in **(H)**. GAPDH used as loading control. Points represent technical replicates from three independent experiments. One-way ANOVA with Tukey’s multiple comparisons correction. **(J)** Both the lactone and salt formulations of 1μM cerivastatin promote STMN2 expression equally via luciferase assay. One-way ANOVA with Tukey’s multiple comparisons correction. Points represent mean of technical replicates from independent experiments. One-way ANOVA with Tukey’s multiple comparisons. **(K)** WT-SH-SY5Y edited to express mGreenLantern on one endogenous STMN2 allele co-cultured with 50% unedited cells exhibit increased endogenous fluorescence upon statin exposure. Points represent individual wells. Unpaired t-test. Scale bar = 50μm. **(L)** Acyl-biotin exchange assay demonstrates no change in relative palmitoylation status of STMN2 after 1μM cerivastatin treatment. Points represent technical replicates from two independent experiments. Unpaired t-test. **(M)** RT-qPCR demonstrating 2μM simvastatin and 1μM cerivastatin treatment returns *STMN2* mRNA expression to WT baseline in TDP-43 mutant (N352S) SH-SY5Y. Points represent mean of technical triplicates from three independent experiments. One-way ANOVA with Dunnett’s multiple comparisons. **(N)** RT-qPCR of statin-treated iPSC demonstrates a dose-dependent increase in *STMN2* RNA expression. One-way ANOVA with Tukey’s multiple comparisons correction. **(O-P)** RT-qPCR **(O)** and immunoblot **(P)** of mouse N2a cells treated with 1μM cerivastatin indicates conservation of the mechanism between species. Points represent averages of technical triplicates from independent biological replicates. One-way ANOVA with Dunnett’s correction.

### Statins promote STMN2 expression independently of TDP-43 function

To confirm that the observed effect was not due to an increase in total cell number, statins were re-assessed for dose-response after normalization of luminescence values using automated counting of nuclei stained with Hoechst **(Fig. 1d; Supplementary Table 3)**. Depending on their side-chain polarity, statins are either lipophilic (e.g cerivastatin, simvastatin, pitavastatin) or hydrophilic (e.g pravastatin, rosuvastatin)^48,49^. Lipophilic statins display broader tissue exposure in non-hepatic tissues, while hydrophilic statins require active-transport across cellular membranes and display higher hepatoselectivity^32,50,51^. Lipophilic, but not hydrophilic statins, increased STMN2 with EC50 in the nanomolar range (**Fig. 1d**; **Supplementary Table 3**). Luciferase assay **(Fig. 1e.)** and immunoblotting **(Fig. 1f.)** confirmed that the lipophilic statins, simvastatin and cerivastatin, returned STMN2 protein expression to WT baseline in TDP-43 mutant lines in a dose-dependent manner.

Statins repress cholesterol synthesis by preventing the production of mevalonate; a substrate also required for the downstream synthesis of the isoprenoids GGPP and FPP^34^. These 20- and 15- carbon moieties are utilized as the basis of the lipid post-translational modifications geranylgeranylation and farnesylation, respectively. To determine target engagement and efficient inhibition of HMG-CoA reductase by statins, we used an electromobility shift assay for HDJ2 (*DNAJA1*), a heat-shock protein farnesylated at its C-terminal^52^. Indeed, inhibition of HDJ2 prenylation can be visualized as an accumulation of the unprenylated form and upward shift in molecular weight after immunoblotting **(Fig. 1g; Extended data Fig. 1j.)**. Cerivastatin and pitavastatin - but not pravastatin - inhibited the farnesylation of HDJ2; demonstrating that the failure of pravastatin to increase STMN2 results from its inability to inhibit HMG-CoA reductase in SH-SY5Y cells.

We next sought to determine whether statins modulate STMN2 expression in the context of TDP-43 depletion. We treated SH-SY5Y cells with siRNA targeting TDP-43, resulting in ∼90% reduction in TDP-43 expression and subsequent STMN2 loss **(Fig. 1h, 1i.)**. Statin treatment elicited a 2-fold increase in STMN2 protein despite TDP-43 depletion as demonstrated by luciferase assay **(Fig. 1h.)** and immunoblotting **(Fig. 1i.)**.

Notably, statin treatment also promoted STMN2 expression in WT cells as shown by luciferase assay **(Fig. 1j; Extended data Fig. 1j.)**, with both the salt and prodrug lactone forms of cerivastatin increasing STMN2 expression equally, likely due to the presence of esterases capable of converting the prodrug in serum-containing media **(Fig. 1j.)**. Confocal microscopy using a CRISPR-edited WT SH-SY5Y line expressing endogenous STMN2 fused with mGreenLantern (*STMN2*-mGL) also demonstrated increased STMN2 expression upon statin treatment **(Fig. 1k.; Extended data Fig. 1k and 1l.)**. Indeed, cerivastatin increased both cytoplasmic and neuritic intensity of endogenous *STMN2*-mGL **(Fig. 1k.)**.

### Statins restore STMN2 levels in TDP-43 deficient cells by promoting its transcription

We next asked whether the observed statin-mediated effect on STMN2 was driven transcriptionally or at the post-translational level. STMN2 is post-translationally modified at its N-terminal membrane attachment motif via a lipid-moiety binding region consisting of two palmitoylated cysteine residues (amino acids 22 and 24) **(Extended data Fig. 1m)** which are essential for the correct subcellular targeting of the protein. Mutation of Cys24 results in abolishment of neurite staining and relocalization of STMN2 to the mitochondria^53–55^. To determine whether the phenotypic effect of statin treatment was occurring via disruption of palmitoylation, we performed acyl-biotin exchange as described previously^56,57^. Lysates were treated with either Tris or Hydroxylamine to cleave thioesters, followed by biotinylation of free thiols and neutravidin extraction of biotinylated proteins prior to immunoblotting of STMN2 protein. Statin treatment did not influence the relative proportion of palmitoylated STMN2 **(Fig. 1l.)**, indicating its effect is not derived from the alteration of N-terminal palmitoylation of Cys22 and Cys24.

Instead, statin treatment promoted transcriptional regulation of STMN2 leading to increased *STMN2* RNA levels in TDP-43 mutant SH-SY5Y **(Fig. 1m.)**. Notably, expression of the *STMN2* cryptic exon in TDP-43 (N352S) SH-SY5Y was proportionately increased relative to the full-length transcript, indicating an overall transcriptional activation regardless (and independent) of TDP-43-induced *STMN2* mis-splicing **(Extended data Fig. 1n.)**. Statins also significantly increased *STMN2* RNA levels in WT induced pluripotent stem cells (iPSC) **(Fig. 1n.)**. Notably, statins also promoted expression of *Stmn2* RNA **(Fig. 1o.)** and protein **(Fig. 1p.)** in murine N2a cells, indicating conservation of the mechanism across species. Together, these results identify statins as potent modulators of STMN2 and indicate their potential to increase STMN2 levels in the context of TDP-43 loss-of-function.

### Statin-mediated increase in STMN2 requires engagement of HMG-CoA reductase

Statins inhibit the production of mevalonate by binding the catalytic domain of HMG-CoA reductase, the rate-limiting enzyme in cholesterol synthesis **(Fig. 2a.)**^31^, however their overall clinical benefit is believed to derive from additional pleiotropic effects^33,58–61^. To confirm that the statin-mediated effect on STMN2 occurs via inhibition of the canonical mevalonate pathway, we first co-treated SH-SY5Y cells with both cerivastatin and downstream substrates. Co-treatment with mevalonate or geranylgeranylpyrophosphate (GGPP), abrogated the statin-mediated increase in STMN2 measured by luciferase assay **(Fig. 2b.)**, immunoblot **(Fig. 2c.)**, and immunofluorescence **(Fig. 2d.)**. Cholesterol supplementation with or without statin co-administration had no impact on STMN2 expression at the protein level **(Extended data Fig. 2a.)**, and minimal effect at the RNA level **(Extended data Fig. 2b.)**. To further confirm that engagement of HMG-CoA reductase is a pre-requisite for STMN2 modulation by statins, we synthesized an analogue of pitavastatin (MGB-06), which lacks the C_5_-hydroxyl. This modification removes a key hydrogen bond to the HMG-CoA reductase catalytic domain **(Fig. 2e, 2f.)**^31^. As expected, MGB-06 was unable to induce STMN2 expression as determined by luciferase assay **(Fig. 2g.)** or immunoblot **(Fig. 2h.)** and did not inhibit HDJ2 farnesylation **(Fig. 2h.)**.

**Figure 2.**
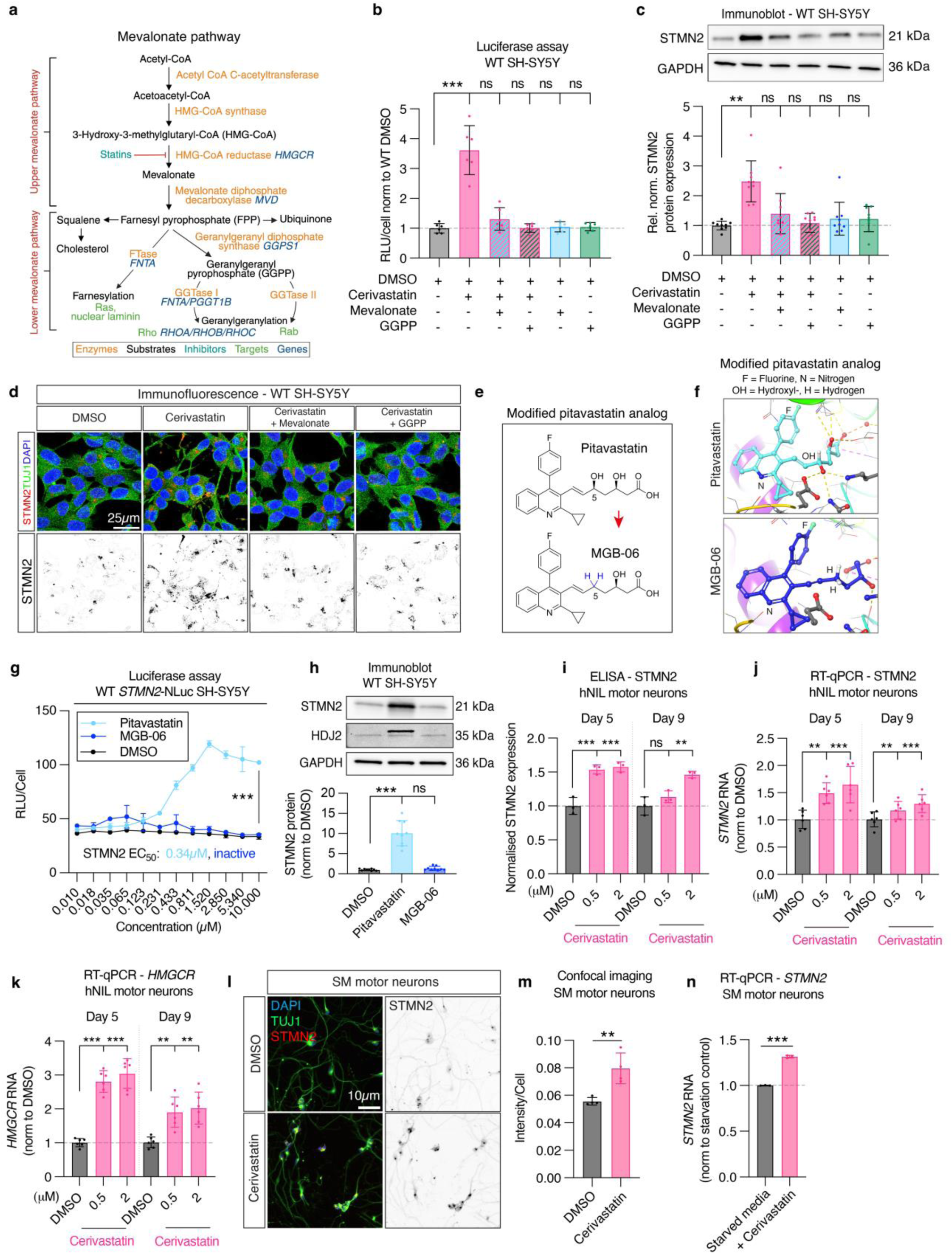
Chemical and genetic inhibition of the mevalonate pathway promotes STMN2 expression. **(A)** Simplified schematic illustrating key steps of the mevalonate pathway. **(B-C)** Luciferase assay **(B)** and immunoblot **(C)** to quantify STMN2 protein in WT SH-SY5Y cells co-treated with 1μM cerivastatin and either 30μM mevalonate or 10μM GGPP substrates. Points represent mean of technical replicates from independent experiments. One-way ANOVA with Dunnett’s multiple comparisons. Asterisks indicate comparison of each condition to DMSO control. **(D)** Immunofluorescence with STMN2 (red/black) and TUJ1 (green) antibodies of WT SH-SY5Y treated with statin and mevalonate/GGPP substrates. Scale bar = 25μm. **(E)** Newly synthesized isoform of pitavastatin (MGB06) bearing modified hydroxyl side-chain to inhibit engagement of the HMG-CoA reductase catalytic domain. **(F)** MGB06 Hydroxyl-modification prevents hydrogen bonding to the HMG-CoA reductase catalytic domain. Panels generated using Maestro software (Schrodinger). (G-H) MGB06 has no effect on STMN2 expression as measured by luciferase assay **(G)** or immunoblot **(H)**. Points represent mean of technical replicates in three independent experiments. Two-way ANOVA with Tukey’s multiple comparisons. Asterisks represent comparisons to DMSO control. **(I-K)** Enzyme-linked immunosorbent assay for STMN2 **(I)** and RT-qPCR for *STMN2* **(J)** and *HMGCR* **(K)** in iPSC-derived hNIL motor neurons treated with cerivastatin and harvested at day 5 or day 9 post-doxycycline induction, normalized to mean of DMSO control values. Points represent replicates from three independent experiments. One-way ANOVA with Holm-Šídak multiple comparisons. Asterisks indicate comparison to DMSO. **(L-M)** Confocal image **(L)** and automated quantification of STMN2 immunostaining **(M)** in motor neurons differentiated from iPSC using a small molecules (SM)-based protocol and treated with cerivastatin. Points represent mean intensity across four independent wells. Unpaired t-test. **(N)** RT-qPCR for *STMN2* in small-molecule differentiated motor neurons. One-way ANOVA with Holm-Šídak multiple comparisons. Asterisks indicate comparison to DMSO.

### Inhibition of the mevalonate pathway promotes STMN2 expression in iPSC-derived neurons

As *STMN2* is neuronally enriched^14^, and in order to demonstrate that statin modulation of STMN2 could be achieved in post-mitotic, differentiated neurons, we used a PiggyBac transposase system for doxycycline-inducible expression of the transcription factors Neurogenin 2 (NGN2), Islet 1, and LIM Homeobox 3 transcription factors and differentiate iPSC into motor neurons (hNIL-motor neurons)^62^. Enzyme-linked immunosorbent assay (ELISA) after cerivastatin treatment exhibited a significant increase in STMN2 protein in hNIL-motor neurons at both day 5 and day 9 post-doxycycline induction **(Fig. 2i.)**. Statin treatment was associated with a significant increase in *STMN2* RNA **(Fig. 2j.)**, as well as a concomitant increase in *HMGCR* RNA **(Fig. 2k.)**, reflecting a feedback response to the inhibition of the mevalonate pathway. Confocal image analysis of iPSC-derived motor neurons differentiated for 21 days using small molecules (SM-motor neurons)^14^ and immunostained with STMN2 antibody demonstrated a similar increase in STMN2 after cerivastatin treatment **(Fig. 2l and 2m.)**, that was also induced transcriptionally **(Fig. 2n.)**, together demonstrating that both the mevalonate pathway and STMN2 can be modulated in iPSC-derived neurons through statin administration.

### Genetic inhibition of geranylgeranylation induces STMN2 expression

Next, we asked whether genetic inhibition of the mevalonate pathway could elicit a similar effect on *STMN2*. As *HMGCR* is the rate-limiting enzyme, and its expression is increased upon statin exposure **(Fig. 2k.)**, we first manipulated its expression in SH-SY5Y using siRNA-mediated repression. Downregulation of *HMGCR* to approximately 50% of normal levels **(Extended data Fig. 2c.)** yielded a significant increase in STMN2 expression measured by luciferase assay **(Fig. 3a)**. In addition, analysis of single-cell RNA-seq following CRISPRi-mediated knockdown of *PGGT1B* - a subunit of GGTase I **(Fig. 2a.)** - in iPSC^63^ showed increase in *STMN2* RNA level **(Fig. 3b.)**. Consistently, treatment of SH-SY5Y using an ASO targeting *PGGT1B* significantly increased *STMN2* RNA levels **(Fig. 3c.)**. These results are consistent with the observation that co-administration of GGPP abrogated the statin-mediated increase in STMN2 **(Fig. 2b-d.)** and support a central role for geranylgeranylation inhibition in the statin-mediated modulation of STMN2 expression. Notably, ASO-mediated reduction of *RHOB* also significantly increased *STMN2* RNA levels **(Fig. 3d.)** suggesting that inhibition of Rho-GTPase prenylation by statin treatment reduces their activity^36,37^ and subsequently promotes *STMN2* expression.

**Figure 3.**
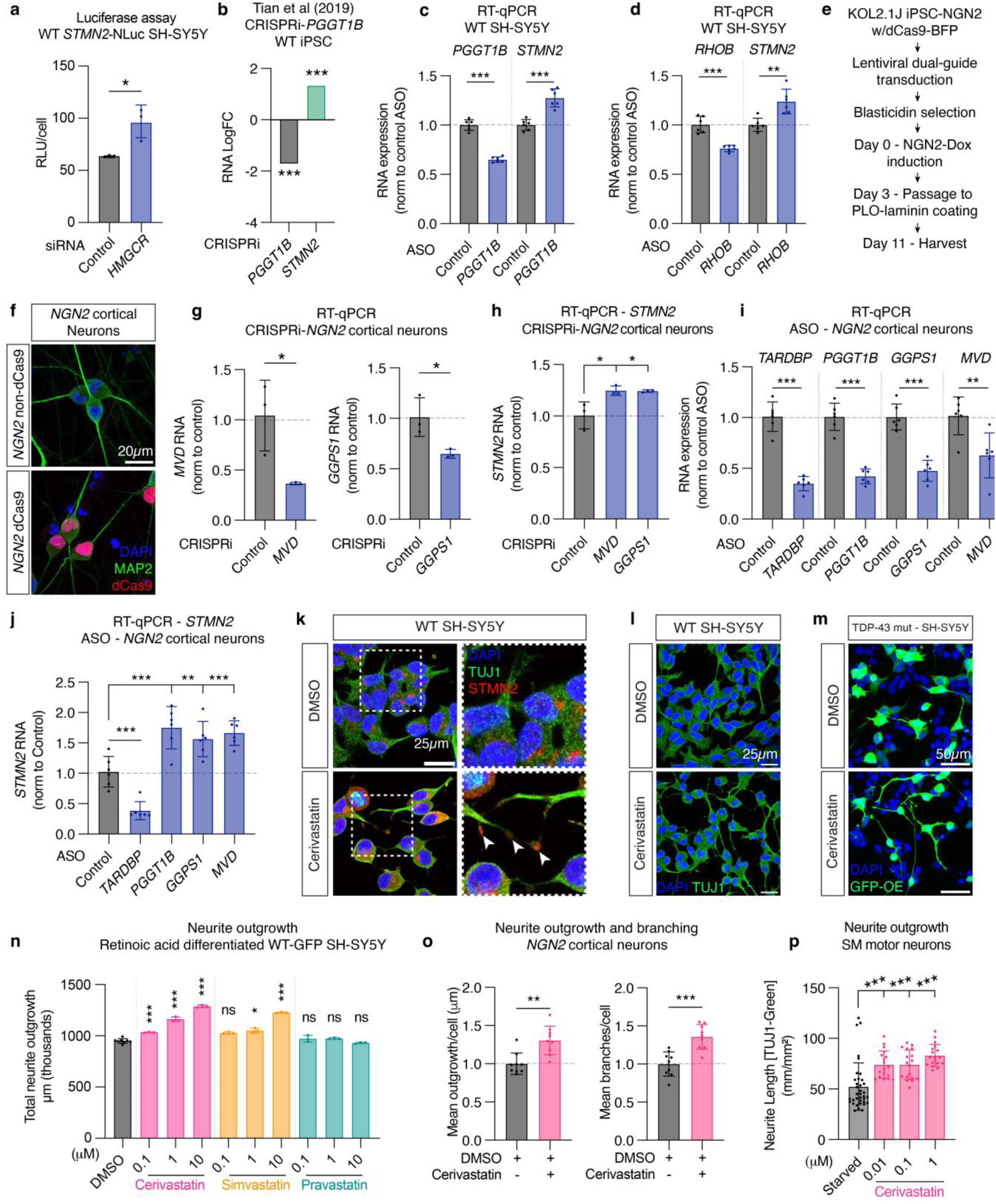
Genetic and pharmacological inhibition of the mevalonate pathway promotes STMN2 expression and neurite outgrowth. **(A)** Luciferase assay for STMN2 in WT SH-SY5Y cells treated with siRNA against *HMGCR* and control. Points represent mean of three technical replicates. Unpaired t-test. **(B)** Pooled CRISPRi-single-cell RNA-seq data from Tian et al (2019)^63^ demonstrates that inhibition of the GGTase I beta-subunit (*PGGT1B)* promotes *STMN2* expression in iPSCs. Asterisks indicate comparison to non-targeting. **(C)** RT-qPCR for *PGGT1B* and *STMN2* transcripts in WT SH-SY5Y cells treated with ASOs targeting *PGGT1B*. Points represent average of technical replicates from independent biological replicates. Unpaired t-tests. **(D)** RT-qPCR for *RHOB* and *STMN2* transcripts in WT SH-SY5Y cells treated with ASOs targeting *RHOB*. Points represent average of technical replicates from independent experiments. Unpaired t-tests. **(E)** Experimental scheme for CRISPRi of targets using NGN2-dCas9 cortical neurons. iPSC were transduced using lentiviral constructs bearing dual-guide RNA at day -3, selected using blasticidin, and NGN2 expression was induced through the addition of doxycycline. Cells were harvested at day 11 post-induction for RT-qPCR. **(F)** Immunofluorescence imaging of *NGN2*-dCas9-BFP iPSC neurons. dCas9 is localized to the nucleus and used to repress indicated genes after lentiviral-mediated expression of dual gRNA. **(G)** CRISPR inhibition of *MVD* and *GGPS1* in iPSC-derived cortical neurons results in a knockdown efficiency of 60% and 40% respectively via RT-qPCR. Points represent average of technical replicates from independent experiments. Unpaired t-tests. **(H)** RT-qPCR for *STMN2* upon CRISPR inhibition of *MVD* or *GGPS1* in iPSC-derived cortical neurons. Points represent average of technical replicates from independent experiments. One-way ANOVA with Dunnett’s multiple comparisons. **(I)** RT-qPCR of selected targets in day 32 iPSC-derived cortical neurons following 10 days of treatment with ASOs targeting *TARDBP, PGGT1B*, *GGPS1* or *MVD*. Points represent independent biological replicates. Unpaired t-tests. (J) RT-qPCR for *STMN2* expression following ASO knockdowns in **(I)**. Points represent independent biological replicates. One-way ANOVA with Dunnett’s multiple comparisons. **(K)** Immunofluorescent staining using STMN2 (red) and TUJ1 (green) antibodies in non-differentiated WT SH-SY5Y cells demonstrate an upregulation in STMN2 expression, including at the nascent growth cone (arrow). Right panels are insets as indicated. Scale bar = 25μm. **(L)** Immunofluorescent staining of WT SH-SY5Y using TUJ1 antibody (green) indicates neurite length extension following statin treatment. Scale bars = 25μm. **(M)** GFP overexpression in TDP-43 mutant SH-SY5Y also reveals neurite extension following 1μM cerivastatin treatment. **(N)** Automated image analysis of retinoic acid differentiated WT-GFP SH-SY5Y demonstrates dose-response increase in neurite length upon treatment with lipophilic cerivastatin and simvastatin, but not hydrophilic pravastatin. Points represent average of technical replicates from independent experiments. One-way ANOVA with Dunnett’s multiple comparisons. **(O)** Image analysis of TUJ1 staining to measure neurite length and branching in *NGN2* iPSC-derived cortical neurons after statin treatment. Unpaired t-tests. Points represent average values from nine images across each well, across two independent experiments. **(P)** Automated quantification of neurite length in iPSC-derived motor neurons treated with cerivastatin and immunostained with a TUJ1 antibody. Points represent average of measurements from individual wells. One-way ANOVA with Dunnett’s multiple comparisons.

We next asked whether manipulation of other genes in the pathway also modulated STMN2. Mevalonate diphosphate decarboxylase (*MVD*) converts mevalonate 5-diphosphate to isopentyl diphosphate, while geranylgeranyl diphosphate synthase I (*GGPS1*) catalyzes the synthesis of geranylgeranyl pyrophosphate (GGPP) from isopentyl diphosphate **(Fig. 2a.)**. Both are essential for producing the GGPP substrate required for prenylation of Rho-GTPases. We engineered iPSCs to allow doxycycline-inducible expression of the NGN2 transcription factor and expression of dCas9 for CRISPRi inhibition of targets in NGN2-cortical neurons **(Fig. 3e, 3f.)**. Doxycycline induced NGN2-iPSC-dCas9 expressed markers of neuronal differentiation including MAP2 **(Fig. 3f.)**, TUJ1, NeuN, and vGlut1 **(Extended data Fig. 2d.)**. Using lentiviral-mediated expression of dual-guide RNAs (gRNAs) against *MVD* or *GGPS1*, we observed a knockdown efficiency of 60% and 40%, respectively, compared to non-targeting dual gRNAs **(Fig. 3g.)**. Genetic inhibition of either *MVD* or *GGPS1* led to ∼25% upregulation in *STMN2* **(Fig. 3h.)**. As an orthogonal approach, we used ASO-mediated downregulation of *PGGT1B*, *GGPS1* and *MVD* in iPSC-derived cortical neurons. ASO treatment from day 22 to day 32 after NGN2 induction resulted in a significant repression of all three genes **(Fig. 3i.)** and a concomitant increase in *STMN2* **(Fig. 3j.)**. Combined, these results demonstrate that both chemical and genetic inhibition of the mevalonate pathway effectively modulates STMN2 in iPSC-derived neurons.

### Statins promote neurite extension

Previous studies have associated increased neurite growth with statin exposure^42–46^. Additionally, CRISPR inhibition of either *HMGCR* or *PGGT1B* results in promotion of neurite extension and branching in iPSC-derived cortical neurons^63^. Since both statin treatment **(Fig. 1 and 2)** and *PGGT1B* repression **(Fig. 3b, 3j.)** resulted in increased *STMN2*, we next asked whether the observed increase in STMN2 was associated phenotypically with increase in neurite length and branching. SH-SY5Y cells demonstrated increased accumulation of STMN2 in nascent growth cones following statin treatment **(Fig. 3k.)**, as well as elongation of neurites in both WT **(Fig. 3l.)** and TDP-43 mutant lines **(Fig. 3m.)**. A significant dose- and statin-dependent increase in neurite outgrowth **(Fig. 3n.)** and branching **(Extended data Fig. 2e.)** was also observed in retinoic-acid differentiated SH-SY5Y. Pravastatin had no effect, consistent with our observation indicating lack of HMG-CoA reductase target engagement in SH-SY5Y cells **(Fig. 1d-g.)**. A similar statin-mediated promotion of neurite outgrowth and branching was observed in *NGN2* iPSC-derived cortical neurons **(Fig. 3o.)** and iPSC-derived motor neurons **(Fig. 3p.)**.

### Statins upregulate cytoskeletal and pro-neuroregenerative transcriptional networks

To identify downstream targets of HMG-CoA reductase inhibition, we conducted RNA-seq of statin-treated SH-SY5Y cells using co-treatment with mevalonate as a negative control **(Supplementary Tables 4, 5, 6)**. Simvastatin was chosen for its potency in screening **(Fig. 1c-f.)**, its clinical availability, and previous use *in vivo*^64,65^. Simvastatin treatment for 24 hours resulted in the significant up/downregulation of 940 and 922 genes, respectively **(Fig. 4a, 4b; Supplementary Table 5)**. Comparison of gene expression in cells treated with simvastatin but supplemented with mevalonate demonstrated correction of almost all transcriptional changes associated with statin treatment **(Fig. 4b.)**. *STMN2* mRNA was itself upregulated ∼1.8-fold, while *TARDBP,* the gene encoding TDP-43, was unchanged **(Fig. 4c.)**, consistent with the statin-mediated effect on STMN2 being unrelated to TDP-43 homeostasis. Another TDP-43-dependent, cryptically mis-spliced gene, *UNC13A*^66,67^, was called as significant by our RNA-seq analysis pipeline, but associated transcript per million (TPM) values were low and variable **(Supplementary Tables 4, 5)**. Subsequent RT-qPCR of cerivastatin treated SH-SY5Y **(Extended data Fig. 3a.)** and iPSC-derived hNIL motor neurons **(Extended data Fig. 3b.)** confirmed a significant upregulation in *UNC13A* mRNA after treatment. Notably, *UNC13B*, a paralog of *UNC13A*, exhibited similarly low TPM but was also significantly upregulated **(Extended data Fig. 3c; Supplementary Tables 4, 5)**.

**Figure 4.**
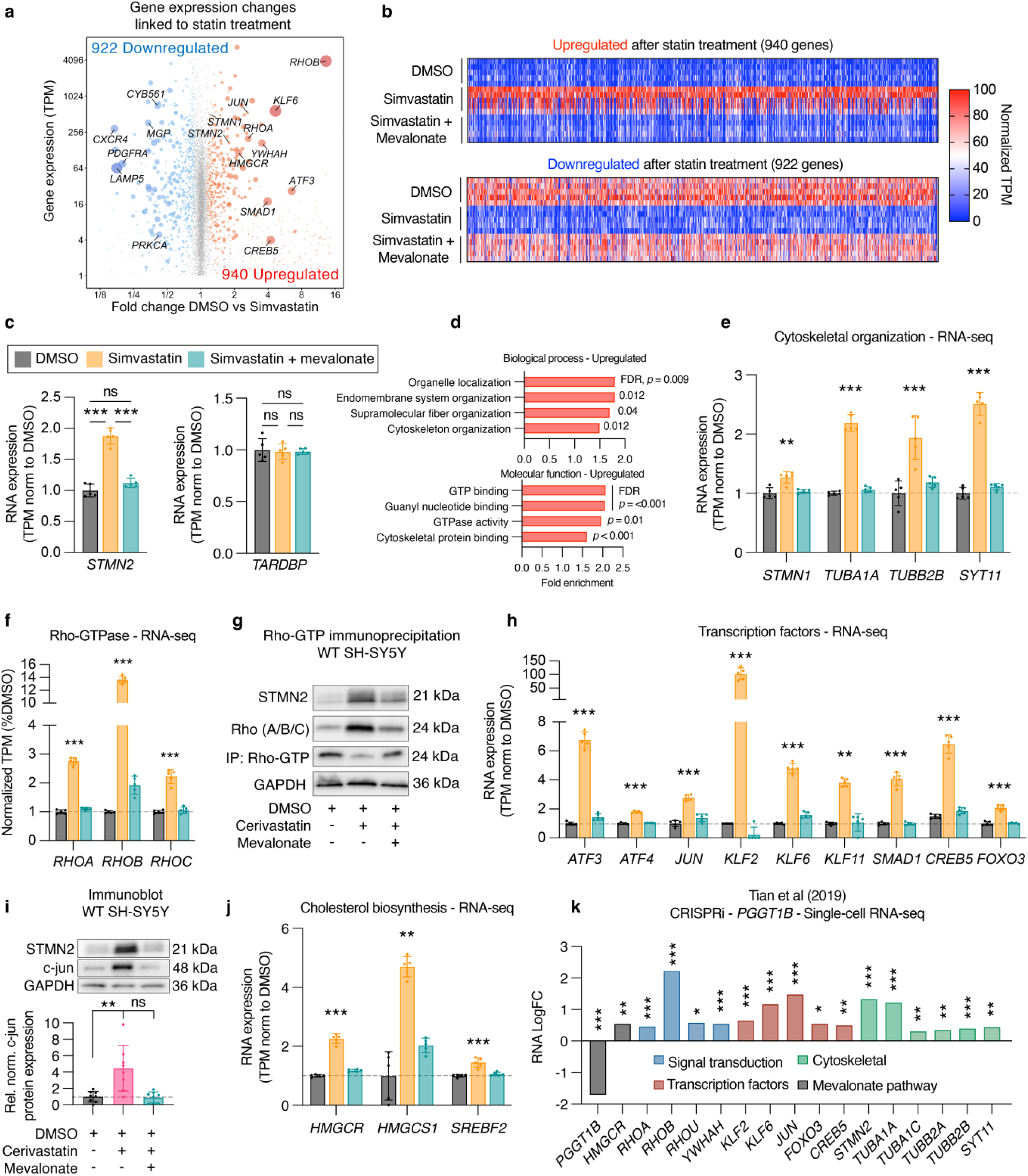
Statins induce the transcription of cytoskeletal, small GTP-ase and regenerative transcription factors. **(A)** Volcano plot from RNAseq analysis of simvastatin or DMSO treated WT SH-SY5Y cells showing significantly downregulated (blue) and upregulated (red) genes. Dot size is inversely proportional to DEseq adjusted p-value. Genes shown: Transcripts Per Million (TPM) 1-15000, Fold change (FC) 0.12-15. **(B)** Heatmaps of normalized TPM of all up/downregulated genes following statin treatment. Samples treated with simvastatin and mevalonate are included as control. Normalization for each point was to mean TPM of DMSO condition. **(C)** TPM normalized to DMSO average for *STMN2* and *TARDBP* transcripts upon simvastatin or simvastatin + mevalonate treatment. One-way ANOVA with Dunnett’s multiple comparisons. **(D)** Top gene ontology terms identified using ShinyGO as enriched in upregulated genes upon statin treatment. (E-F) Normalized TPM for selected cytoskeletal genes **(E)** and Rho-GTPases **(F)**. Asterisks indicate adjusted p-values from comparison between DMSO and Simvastatin treatments using DESeq2. **(G)** Immunoprecipitation using a RhoA/B/C antibody cocktail revealed an upregulation of total Rho, but downregulation GTP-bound Rho, after 1μM cerivastatin treatment. **(H)** Normalized TPM for selected transcription factors. Asterisks indicate adjusted p-values from comparison between DMSO and Simvastatin treatments using DESeq2. **(I)** Immunoblot with antibodies against STMN2 and c-jun in WT SH-SY5Y cells treated with 1μM cerivastatin alone or with the addition of mevalonate. GAPDH is included as loading control. Points represent technical replicates from three independent experiments. One-way ANOVA with Dunnett’s multiple comparisons. **(J)** Normalized TPM for selected cholesterol biosynthesis gene. **(K)** Examples of genes upregulated both after simvastatin treatment and CRISPR inhibition of *PGGT1B*^63^. Asterisks indicate p-value of transcriptional changes in iPSC with *PGGT1B* gRNA compared to control gRNA ^63^.

Gene Ontology (GO) analysis revealed an upregulation of networks related to cytoskeletal organization (e.g *STMN1, STMN2, SYT11, TUBB3, TUBB2B, TUBA1A, NES, ACTN1*; FDR = 0.01) and GTP-ase activity (e.g *RHOA, RHOB, RHOC, CDC42*; FDR <0.001) **(Fig. 4d, 4f.; Supplementary Table 6)**. Downregulated pathways included those enriched in tube development, blood circulation, and angiogenesis **(Extended data Fig. 3d., Supplementary Table 6)**. Several Rho-family genes were upregulated following statin treatment **(Fig. 4f.)**, consistent with a cellular response to the inhibition of their prenylation. Immunoprecipitation using a RhoA/B/C antibody cocktail validated the upregulation of total Rho, but indicated a decrease in the active GTP-bound form **(Fig. 4g; Extended data Fig. 3e.)**. This result suggests that while levels of Rho GTP-ases are increased, inhibition of their prenylation by statin treatment decreases their activity.

In addition, transcription factors reported to promote axon regeneration^68–73^ including *KLF6, KLF2, SMAD1*, CREB5, *JUN*, *ATF3,* and *ATF4* were significantly upregulated **(Fig. 4h.)**. Increase of nuclear c-Jun upon cerivastatin treatment was confirmed using immunofluorescence **(Extended data Fig. 3f.)** and immunoblotting **(Fig. 4i.)**. Notably, c-Jun and ATF3 combine dimerically to form one subclass of the AP-1 complex, and regulate transcription by binding the 12-O-Tetradecanoylphorbol-13-acetate (TPA) response element (TRE) via the palindromic consensus sequence 5’-TGA[G/C]TCA-3’^74^. Predictive analysis indicated partial TRE and cAMP response element (CRE) sites clustered in the proximal *STMN2* promoter **(Extended data Fig. 3g.)**, supporting a role for these statin-induced transcription factors in the regulation of *STMN2* transcription.

Finally, several genes involved in cholesterol/sterol biosynthetic process (e.g *HMGCR, SQLE, IDI1, HMGCS1, SREBF2*; FDR = 0.06) and cellular response to starvation (e.g *FOXO3, ATF3, ATF4*, *SREBF2*; FDR = 0.07) **(Fig. 4j; Supplementary Table 6)** were also upregulated, consistent with a feedback response to cholesterol lowering.

As both chemical and genetic inhibition of geranylgeranylation promotes STMN2 expression **(Fig. 2, 3)**, we next compared our RNA-seq data with single-cell RNA-seq following *PGGT1B* CRISPRi knockdown in iPSCs generated by Tian et al (2019)^63^. The comparison yielded a strikingly similar transcriptional response with 148 genes differentially expressed in both datasets, 106 of them in the same direction **(Supplementary Table 7)**, including a significant upregulation of *STMN2*, *HMGCR*, *SYT11*, *TUBA1A* and *TUBB2B* **(Fig. 3b, 4k.)**. Notably, *RHOB* was the top differentially expressed gene in both datasets, with transcription factors including *FOXO3*, *JUN*, *KLF6* and *KLF2* also increased both upon statin treatment and *PGGT1B* inhibition **(Fig. 4k, Extended data Fig 3h., Supplementary Table 7)**. This comparison highlights a common transcriptional response by statin treatment in SH-SY5Y and *PGGT1B* repression in iPSC, and supports a mechanistic link between Rho isoprenylation and *STMN2* expression.

### ATF3/Atf3 is a conserved regulator of STMN2 expression

Since statin treatment promotes both *STMN2* and *ATF3* expression **(Fig. 1, 2, 4)**, overexpression of *Atf3* elicits neuroprotection in *SOD1*-G93A mice^70,75^, and both *STMN2* and *Atf3* are upregulated in response to neuronal injury^71–73^, we next asked whether ATF3 regulates *STMN2* expression *in vitro* and *in vivo*. First, we validated that induction of *ATF3* expression by statin treatment also occurred in differentiated neurons, including ReNVM-derived neurons^76^ **(Fig. 5a.)** and hNIL-motor neurons **(Fig. 5b.)**. To assess the impact of ATF3 ablation *in vitro,* we then generated an *ATF3* knockout SH-SY5Y line using CRISPR-Cas9 **(Extended data Fig. 4a, 4b**). Genome editing using a single gRNA targeting exon 1 resulted in a homozygous single base-pair deletion, leading to insertion of a premature stop codon and non-sense mediated decay as demonstrated by reduced *ATF3* RNA levels compared to a control line generated with a non-targeting gRNA **(Fig. 5c)**. Expression of the known ATF3 targets *UTP23*, *TRIB3*, *SOX9* and *SPRY1*^77,78^ were significantly reduced in *ATF3* KO cells **(Extended data Fig. 4c)**. Both RT-qPCR **(Fig. 5d.)** and ELISA **(Fig. 5e.)** demonstrated that the statin-mediated STMN2 response was partially repressed in *ATF3* KO cells, indicating a direct role for ATF3 in the induction of STMN2 expression upon statin treatment.

**Figure 5.**
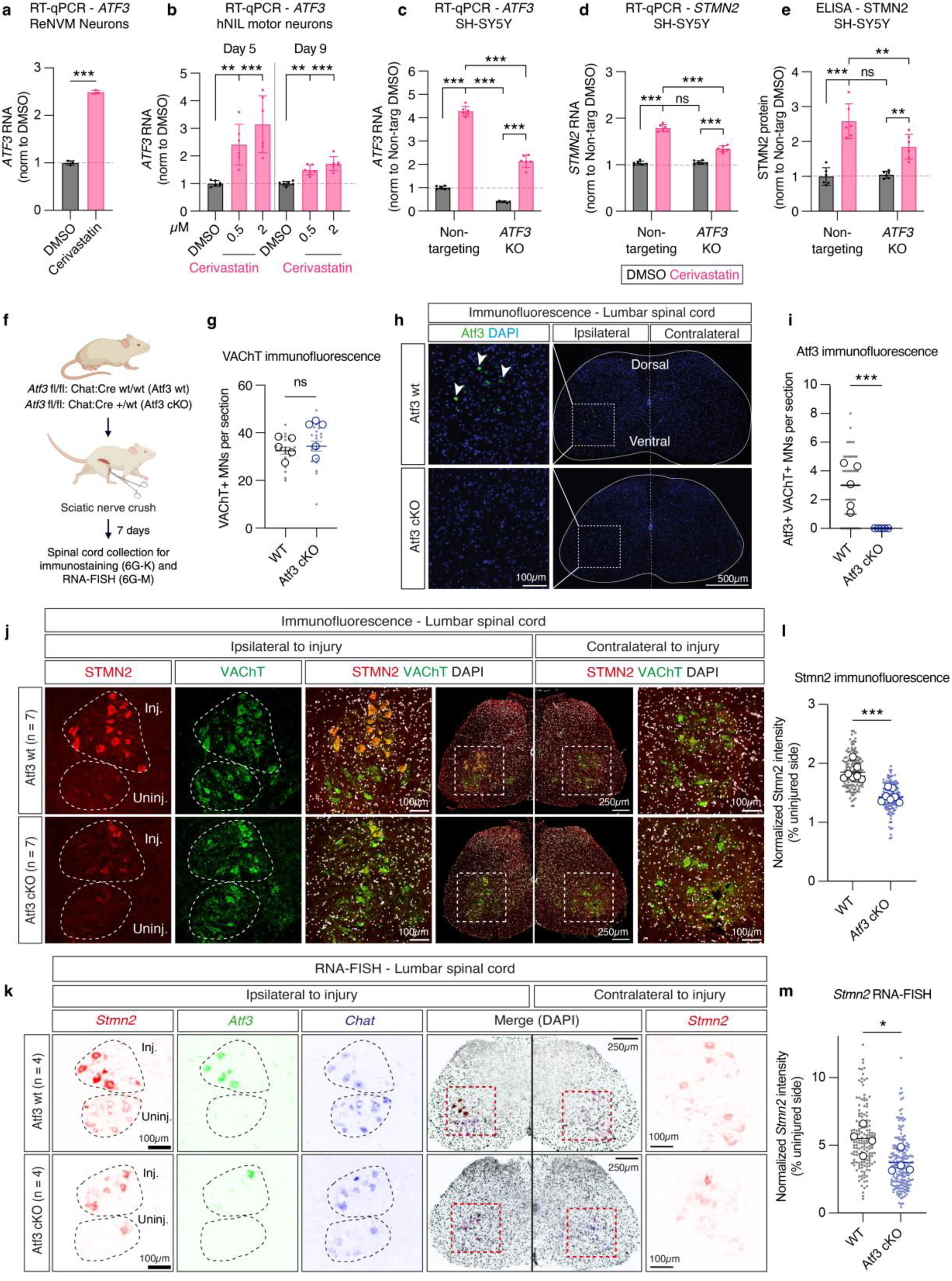
ATF3 is a conserved regulator of STMN2 expression. **(A)** RT-qPCR for *ATF3* in differentiated ReNVM neurons treated with DMSO or 2μM cerivastatin. Points represent the mean of technical replicates from three independent experiments. Unpaired t-test. **(B)** RT-qPCR for *ATF3* in hNIL motor neurons treated with DMSO or 0.5 and 2μM cerivastatin at day 5 or day 9 of differentiation. Points represent average of technical replicates from three independent experiments. One-way ANOVA with Dunnett’s multiple comparisons. **(C)** RT-qPCR for *ATF3* in SH-SY5Y CRISPR-edited to inactivate ATF3 compared to SH-SY5Y cells treated with a non-targeting sgRNA. Points represent mean of technical replicates from independent biological replicates. Two-way ANOVA with Tukey’s multiple comparisons correction. **(D-E)** RT-qPCR **(D)** and ELISA **(E)** for STMN2 RNA and protein, respectively, in ATF3 KO and control lines after 1μM cerivastatin treatment. Points represent average of technical replicates from independent biological replicates. Two-way ANOVA with multiple comparisons Tukey’s correction. **(F)** Schematic of in vivo experimental design in Atf3 conditional knockout mice where expression was removed from motor neurons using Chat-Cre expression. Control (n=7) and Atf3 cKO mice (n=7) underwent surgical sciatic nerve crush and were euthanized 7 days post-injury for tissue analyses. **(G)** Quantification of VAChT+ motor neurons in the lumbar spinal cord. Large points = individual animals, small points = individual neurons. Error bars = mean with SEM. Unpaired t-test. **(H)** Immunofluorescence of the lumbar spinal cord with an Atf3 antibody demonstrates increase of Atf3 expression in the injured motor neurons of control animals but not Atf3 cKO mice. Scale bar inset = 100μm, zoom out = 500μm. **(I)** Quantification of immunofluorescence staining for Atf3 from confocal microscopy shown in **(H)**. Large points = individual animals, small points = individual neurons. Error bars = mean with SEM. Unpaired t-test. **(J)** Immunofluorescent staining with STMN2 and VAChT antibodies of sections from the lumbar spinal cord from control and *Atf3* cKO mice after sciatic nerve crush. Injured and uninjured pools of motor neurons are labelled on the ipsilateral injured side. Scale bars on insets = 100μm, whole spinal cord section = 250μm. Insets of the ventral horns ipsilateral and contralateral to the sciatic nerve injury are highlighted by white dashed boxes on the whole spinal cord sections. **(K)** RNA-FISH of lumbar spinal cord with probes against *Stmn2*, *Atf3* and *Chat* of sections from the lumbar spinal cord from control and *Atf3* cKO mice after sciatic nerve crush. Injured and uninjured pools of motor neurons are labelled on the ipsilateral injured side. Scale bar on insets = 100μm, whole spinal cord section = 250μm. Insets of the ventral horns ipsilateral and contralateral to the sciatic nerve injury are highlighted by red dashed boxes on the whole spinal cord sections. **(L)** Quantification of STMN2 protein levels from immunofluorescence staining in **(J)**. Values of STMN2 intensity in injured motor neurons are normalized to uninjured motor neurons. Small points = individual neurons, large points = individual animals (n=7/condition). Error bar = median. Unpaired t-test. **(M)** Quantification of *Stmn2* RNA levels from FISH staining in **(K)**. Values of *Stmn2* intensity in injured motor neurons are normalized to uninjured motor neurons. Small points = individual neurons, large points = individual animals (n=4/condition). Error bar = median. Unpaired t-test.

Finally, to assess the effect of *Atf3* ablation *in vivo*, we generated a motor-neuron specific *Atf3* conditional knock-out (*Atf3* cKO) mouse model whereby *Atf3* is flanked with LoxP sites (Atf3^fl/fl^) and expression removed from motor neurons via expression of Chat-cre recombinase. Since previous studies have indicated that Atf3 expression is upregulated upon axonal injury^69,72,79^, we performed surgical sciatic nerve crush in *Atf3* cKO and *Atf3* wt mice and collected lumbar spinal cords seven days post injury for immunofluorescence and RNA fluorescence *in situ* hybridization (RNA-FISH) **(Fig. 5f.)**. While the number of cholinergic (VAChT+) spinal motor neurons was similar in *Atf3* wt and *Atf3* cKO mice **(Fig. 5g.)**, Atf3 expression was only induced in injured motor neurons of Atf3 wt mice **(Fig. 5h, 5i.)**. Immunostaining **(Fig. 5j.)** and RNA-FISH **(Fig. 5; Extended data Fig. 4d.)** revealed a significant increase in Stmn2 protein and RNA levels in injured motor neurons of both *Atf3* wt and *Atf3* cKO mice. However, this Stmn2 protein/RNA upregulation in the injured pool of ventral horn neurons was partially alleviated in *Atf3* cKO mice **(Fig. 5j-m)**, indicating that Atf3 regulates the Stmn2 response to axonal injury *in vivo*.

## Discussion

Reduced expression of STMN2 has emerged as a pathological hallmark and an attractive therapeutic target in several neurodegenerative diseases associated with TDP-43 proteinopathy including ALS, FTD, and AD^14–16,18,20^. STMN2 is crucial for axonal regeneration and the maintenance of synaptic connectivity^14,26,80–82^. Increased STMN2 levels elicits neurite outgrowth^25,83^, while loss of Stmn2 in mice is sufficient to produce progressive motor and sensory deficits, including neuromuscular junction denervation and axonal collapse^27–29^. Here, we identify statins as potent inducers of STMN2 expression independently of TDP-43 function. While statins do not block mis-splicing of *STMN2* pre-mRNA, they increase its transcription and restore STMN2 protein level in cells with mutant or depleted TDP-43. Indeed, statin-mediated inhibition of HMG-CoA reductase, a central enzyme in the mevalonate pathway, elicits transcriptional upregulation of STMN2 and other cytoskeletal proteins, while promoting neurite outgrowth. By synthetizing a chemical isoform of pitavastatin unable to bind the HMG-CoA reductase catalytic domain, we confirmed that engagement of HMG-CoA reductase is a pre-requisite for the statin-mediated modulation of STMN2. Using a target engagement assay based on HDJ2 farnesylation, we further demonstrated that hydrophilic statins (rosuvastatin, pravastatin) which failed to alter STMN2 levels in our screen, were, in fact, ineffective at inhibiting the mevalonate pathway.

Statins inhibit the production of mevalonate by binding the catalytic domain of HMG-CoA reductase, the rate-limiting enzyme in cholesterol synthesis^30,31^, however their overall clinical benefit is believed to derive from additional pleiotropic effects^58^. Indeed, mevalonate is also required for the synthesis of geranylgeranyl pyrophosphate (GGPP) and farnesyl pyrophosphate (FPP)^34^ which are used for the post-translational prenylation (geranylgeranylation and farnesylation, respectively) of essential proteins such as Ras, Rho and Rab.

While the regulation of STMN2 by statins is a novel finding, prior studies demonstrated a pro-regenerative effect of statins both *in vitro*^42–46^ and *in vivo*^42,47,84,85^. Indeed, statins were the top hits in a phenotypic screen for neurite growth of stem cell-derived motor neurons, conferring the ability to overcome the neurite inhibitory effect of matrix-associated glycoprotein^42^. Statin treatment was also shown to promote axonal recovery after optic nerve^42^ and sciatic nerve crush^84,85^ in rats. Importantly, statin treatment slowed motor neuron degeneration and improved motor phenotypes in both wobbler^47^ and *SOD1* G93A mice^86^.

In addition, while epidemiological studies yielded contradictory results^87,88^, an increasing body of work suggests that statin usage is associated with lowered disease risk or improved prognosis in both dementia^89–94^ and ALS^86,95–97^. Notably, protein prenylation, a post-translational modification inhibited by statins, was found to be increased in postmortem tissues from patients with ALS ^42^ and AD^98,99^, with evidence suggesting a beneficial effect when reducing/inhibiting prenylation enzymes in AD mouse models^100–104^. Although not well characterized, statins and Farnesyltransferase inhibitors may reduce accumulation of protein aggregates, presumably through lysosomal activation and the proteolysis of disease-specific toxic proteins^102,103,105–108^. Thus, inhibition of the mevalonate pathway may confer benefits across neurodegenerative diseases irrespective of TDP-43 pathology. However, the mechanisms by which statins exert neuroprotection remain largely unknown.

Here, we provide evidence that the statin-mediated effect on STMN2 arises through the inhibition of geranylgeranylation. Indeed, co-administration of GGPP with cerivastatin abrogated the statin-induced increase in STMN2 level. In addition, we found that reduction of the GGTase I catalytic subunit *PGGT1B,* which was previously shown to promote neurite outgrowth^63^, also increases STMN2 expression. Conversely, elevated *PGGT1B* expression in laser-captured motor neurons from sporadic ALS patients correlates with an earlier disease onset^42^. Both GGTase I and FTase act upon CaaX motifs, where ‘C’ is the prenylated cysteine, ‘a’ are aliphatic amino acids, and X determines the catalyzing enzyme^109^. However, this motif is not present on STMN2, indicating the effect arises instead from altered activity of prenylation-dependent signaling proteins. Interestingly, we found that statin treatment elicits a similar transcriptional profile as *PGGT1B* CRISPRi-mediated knockdown^63^, including increased expression of cytoskeletal proteins, pro-regenerative transcription factors^68^ and Rho-GTPases, with RHOB being the top differentially expressed gene in both datasets. Rho GTPases require prenylation for membrane localization and activation^110,111^; thus while statin treatment elevated RhoA/B/C expression, it diminished the fraction of their active GTP-bound form. Notably, RhoA is strongly inhibitory to neuritic extension through its effect on actin polymerization and growth cone collapse^40,41^. Together, this suggests that statins promote STMN2 and neurite extension by inhibiting the prenylation of Rho-GTPases via prevention of their membrane localization and repression of their activity. We note that hyperactivation of RhoA and its downstream effector ROCK has been documented in neurodegenerative diseases such as ALS, where ROCK inhibition has shown benefit in preclinical studies and promising outcomes in clinical trials^111–114^.

The regulatory mechanisms controlling *STMN2* are poorly understood, and uncovering transcription factors that promote its expression is critical. We show that statin treatment increases levels of transcription factors known to promote axonal regeneration^68^. The AP1 complex subunit ATF3 was of particular interest because of its peripheral regenerative effect^115^ and recent work demonstrating its protective effect in *SOD1*-ALS mice^70,75^. CRISPR-Cas9–mediated ATF3 inactivation in cells, together with Cre-dependent conditional *Atf3* knockout in mice, revealed that ATF3 is required for transcriptional activation of the STMN2 gene following statin treatment and axonal injury. These findings demonstrate that STMN2 expression is dynamically regulated at the transcriptional level by a lipid-modification–dependent signaling axis that promotes axonal regenerative capacity.

Overall, this study uncovers mechanistic insights into STMN2 regulation and highlights pharmacological and genetic inhibition of the mevalonate pathway as a promising therapeutic avenue for neurodegenerative diseases.

## Materials and Methods

### SH-SY5Y, RenVM, and iPSC-derived motor neuron cell culture

Neuroblastoma cells (SH-SY5Y) were cultured in DMEM/F12 (Gibco) supplemented with 10% fetal-bovine serum (Omega) and 1% penicillin-streptomycin (Gibco) at 37 °C with 5% CO2. Where indicated, SH-SY5Y were differentiated using 10μM retinoic acid and 50ng/ml BDNF. Homozygous TDP-43 mutant (N352S) SH-SY5Y were generated as previously described^14^. Mouse N2a cells were cultured in DMEM (Gibco) supplemented with 10% fetal-bovine serum (Gibco) and 1% penicillin-streptomycin (Gibco).

ReNcell VM human neural progenitors (Millipore #SCC008)^76,116^ were cultured on Matrigel-coated plates in Dulbecco’s Modified Eagle Medium/F12 media supplemented with 2% B27 neural supplement (ThermoFisher Scientific, #175004044), 20 µg/mL human EGF (Sigma-Aldrich, #E9644), 25 µg/mL bFGF (Stemgent, #03-0002), 0.2% heparin solution (StemCell Technologies, #07980), and 1% Penicillin-Streptomycin. ReNcell VM neural progenitors were differentiated into neurons by removing the growth factors (hEGF and bFGF) from the culture medium. Cells underwent differentiation for 11 days with half-media supplementation every 2 days.

For small molecule iPSC-derived motor neuron differentiation, undifferentiated induced pluripotent stem cells (iPSC) were cultured in StemFlex medium (Gibco) with 1X StemFlex supplement (Gibco) on vitronectin coated plates. Differentiation and culture was performed externally (iXCells, San Diego, USA). A sporadic ALS (sALS) patient iPSC line (40HU-006) was differentiated into motor neurons (SM-motor neurons) using a proprietary small-molecule differentiation protocol patented by iXCells (Medium A, MD-0022A) with growth-factor restriction. Cells were differentiated to day 21 and maintained for a further 7 days, after which either 0.1% DMSO or cerivastatin was added. For RT-qPCR, cells were harvested after 24hrs treatment. For TUJ1 staining and neurite analysis, cells were treated for 72hrs. All compound treatments in other experiments were for 24hrs unless otherwise indicated.

For iPSC-hNIL motor neuron culture, iPSC bearing the hNIL construct were differentiated as previously described^62,117^. Briefly, 90% confluency iPSCs in a 6-well plate were collected and nucleofected with equal concentrations of PiggyBac hNIL donor. To reduce silencing of long cDNAs^118^. we introduced the second intron of HBB into the NIL sequence of Addgene #197089 and PiggyBac transposase plasmids (Addgene). The hNIL construct is designed to allow the expression of the transcription factors *NGN2*, *ISL1* and *LHX3* in a Tet-ON system; it also includes a BFP reporter and puromycin resistance. Two days after nucleofection, medium was changed to iPSC medium + 5ug/mL puromycin to allow selection of nucleofected cells. Nucleofected iPSCs were then expanded to confluence and replated (Day 0) in vitronectin-coated T75 flasks with induction medium (97% DMEM/F12 + 1% N2, 1% NEEA, 1% GlutaMax, 2 ug/mL doxycycline, and 0.1 ug/ mL compound E). After 3 days, cells were collected and replated in poly-L-ornithine (Sigma) coated plates (6 well-plates for protein and RNA collection, glass-bottom 96 well plates for imaging, day 3) with differentiation medium (Neurobasal medium + 1% N2, 1% NEEA, 1% GlutaMax, 0.2% BME, 1 ng/mL BDNF, 1 ng/mL GDNF, 1 ng/mL IGF-1, 1 ng/mL CNTF, 10 nM retinoic acid, 2 ug/mL doxycycline, and 0.1 ug/ mL compound E). Half media changes were performed every 2 days. Statin treatments prior to RT-qPCR and ELISA were for 24 hrs.

For *NGN2* iPSC cortical neuron differentiation, iPSC bearing the NGN2 overexpression system were developed and cultured as previously described^62^. Induction was performed as hNIL but without the addition of compound E. Long-term NGN2 culture media was comprised of Neurobasal medium (Gibco, 21103-049) + 1% N2 supplement (Gibco, 17502-048), 2% B27 supplement (Gibco, #17504-044), 1% GlutaMax (Gibco, 35050061), 1ng/mL BDNF (Life Technologies, PHC7074), 1ng/ml NT3 (Peprotech, #450-03), 50ng/ml Aphidicolin (CellSignaling, #32774) and 2μg/mL doxycycline (Sigma-Aldrich, D5207). Half medium changes were performed every 3 days. For both systems, 50nM Chroman I (MedChem Express, #HY15392) and Emricasan (Selleckchem, #S7775) were used when re-plating and removed after 24 hours. Compound treatments in *NGN2* iPSC-derived cortical neurons were for 48hrs.

### Generation of STMN2-NLuc and STMN2-mGreenLantern SH-SY5Y lines

WT and TDP-mut (N352S) SH-SY5Y cells were edited using CRISPR-SpCas9 nuclease and homology directed repair (HDR) to fuse Nanoluciferase at the C-terminus of both endogenous *STMN2* alleles **(Extended data Fig. 1a.)**. A single guide RNA (sgRNA) targeting *STMN2* was designed (Benchling webtool) and cloned into pSpCas9–2A-green fluorescent protein (GFP) plasmid (px458-Addgene) using the BbsI restriction site. The spacer sequence of the sgRNA targeting *STMN2* was: TGTCTGGCTGAAGCAAGGGA **(Supplementary Table 8)**. For the HDR template, overlapping fragments of 800bp *STMN2* homology arms flanking Nanoluciferase (NLuc, Promega) were generated and ligated into pUC57 backbone. A 42bp GS-linker was also included 5’ to NLuc to prevent steric hindrance of the reporter to the final *STMN2* exon **(Extended data Fig. 1a.)**. Fragments were ligated using Gibson Assembly and nucleofected simultaneously with pSpCas9–2A-GFP-gRNA plasmid using the Amaxa Nucleofector (assay A-033). Forty-eight hours following electroporation, cells were collected, and GFP-positive cells sorted. After expansion, cells were single-cell seeded into 96-well plates using a SH800S Sony cell sorter. Individual clones were expanded, and DNA was extracted for PCR amplification of the *STMN2* Ex5 genomic locus using primers within NLuc and outside the *STMN2* 5’ homology arm to confirm presence of the insert **(Extended data Fig. 1b.)**. The entire *STMN2* coding region and all exon–intron junctions (>500-bp length of each intronic region) were then sequenced in selected homozygous clones to verify the absence of any additional DNA alteration. Primer configurations spanning the entire edited region revealed no CRISPR-induced concatemer formations. The same process and gRNA were used to tag a single endogenous *STMN2* allele with mGreenLantern **(Extended data Fig. 1k, 1.)**. All RT-qPCR and western blotting was performed with isogenic SH-SY5Y cell lines that had undergone the mutagenesis attempt but not acquired the insertion during the original screening process. A full list of genotyping primers used is included in **Supplementary Table 9**.

### Small-molecule screening and validation using Nanoluciferase assay

TDP-mut *STMN2*-NLuc SH-SY5Y cells were used in small-molecule screening at ICCB-Longwood Screen Facility, Boston, MA. 3000 cells/well were plated in white 384-well plates using a 50ul sample volume. 24 hours after plating, compounds were added using 100nl pin transfer. Treatment duration was 24 hours. 25ul NLuc assay was then added to each well according to manufacturer’s instructions (Promega, Cat #N1130). Plates were shaken for 1 min prior to luminescence readings using EnVision2 machine (Perkin Elmer). DMSO-only and WT *STMN2*-NLuc cells were included as negative and positive controls respectively on each plate, producing a median Z-prime factor of 0.53 across the screen. Full screening results of 11,985 unique compounds are included in **Supplementary Table 1**. Libraries were selected based on their relevance to neurodegeneration and known mechanism of action. Each library plate was conducted in duplicate, and Z-scores calculated for each compound. Duplicates were highly correlated **(Extended data Fig. 1h.)**. Z-score >3.0 was used to initially designate hits, and from these, 52 compounds were selected for dose-response analysis (cherry picking) **(Supplementary Table 2)** using a 12-step dilution series ranging 0.03-83μM. For subsequent validation experiments, cell numbers in each well were quantified after treatment by hoescht staining at 1ug/ml for 15 mins at 37C in black, clear bottom plates, prior to imaging at 4X using automated microscopy (ImageXpressMicro). Nuclei were then counted using a custom MATLAB algorithm (https://github.com/waingerlab/Nolan_etal) and luciferase assay run as above. Luminescence results were then normalised against cell counts as RLU/cell. A full list of compounds used for subsequent validation experiments is included in **Supplementary Table 8.**

### Short-interfering RNA (siRNA) and antisense-oligonucleotide (ASO) treatments

For siRNA treatment, cells were reverse transfected with SMARTpool ON-TARGETplus siRNA or control siRNA pool (D001810–10) (GE Dharmacon) at a final concentration of 50 nM, for 72 h after complexing with Lipofectamine RNAiMAX (Invitrogen) in Opti-MEM (Gibco) for 10 min. siRNA was re-dosed at 48hrs. For luciferase assay, cells were then stained with hoescht and imaged 4X using automated microscopy (ImageXpressMicro) prior to Luciferase assay as above. For ASO treatment in SH-SY5Y, cells were lipofected using Lipofectamine 3000 (Thermo) according to manufacturer’s instructions for 48hrs prior to harvesting. For ASO treatment in iPSC cortical neurons, 5μM treatments were begun at day 22 post-doxycycline addition, and re-dosed every three days at each media change. Treatment was performed using free uptake of ASO in media. Total treatment length was 10 days. A full list of the siRNA and ASO used is included in **Supplementary Table 8.**

### Neurite extension analysis

Neurite extension analysis was conducted externally (TCG Life Sciences, India; iXCells, San Diego, USA). WT-GFP SH-SY5Y cells were plated in 96-well, laminin-coated black, glass-bottom plates. Day 2 onwards, 10μM retinoic acid was added. Day 6, 50ng/ml BDNF was added. Day 8, test compounds were added. 24hrs later image acquisition was performed using ImageXpress automated cell imaging system (Molecular Devices). 20X images tiling the totality of each well were stitched and analysed using CellReporterXpress acquisition and analysis software (Molecular Devices), with an n=3 for each condition. To detect neuronal morphological changes in sALS patient-derived iPSC motor neurons, cells were fixed using 4% paraformaldehyde and immunostained using a TUJ1 primary antibody. TUJ1-positive motor neurons were imaged using a BioTek Cytation 5 Plate Reader at 20X magnification. Neurites were automatically traced as previously described using NeurphologyJ implemented as a plugin to ImageJ. Four fields were quantified in each well, with an n=4 wells for each condition. For *NGN2* cortical neuron analysis, images for quantification were obtained at 10X Plan Apo objective using the ImageXpress Confocal high-content imaging microscope (Molecular Devices). Measurements were quantified using MetaXpress® software (Molecular Devices) using the <NEURITE Outgrowth> module. A TUJ1/MAP2 mask was used for neuron cell body and outgrowth detection, and a Hoechst nuclear mask was used for assisting cell body identification. Mean outgrowth per cell measured the average skeletonized outgrowth in μm corrected for diagonal lengths divided by the number of cells. Mean branches per cell measured the total number of branching junctions divided by number of cells.

### Immunoblotting

Total-cells extracts were collected in radioimmunoprecipitation lysis buffer (RIPA buffer). Proteins concentrations were determined by BCA assay (Thermo, cat# 23225), and samples boiled for 10 min in Laemlli buffer with beta-mercaptoethanol prior to running in 4-20% Tris-HCL gels (Bio-rad). Proteins were transferred to PVDF membrane using iBlot transfer (Invitrogen), before blocking in 5% milk solution in tris-buffered saline and 0.1% Tween-20 (TBST) for 1hr and overnight incubation with primary antibodies listed in **Supplementary Table 8**. Immunoblots were washed in TBST and probed with horseradish peroxidase-conjugated secondary antibodies diluted 1:10,000 for 2hr at room temperature (Abcam), before being imaged using Chemidoc MP (Biorad). Target engagement within the mevalonate pathway was assessed using electromobility-shift assay (EMSA) as previously described ^52^. Lysates were run as above and blots stained using anti-HDJ-2 antibody (MA5-12748, ThermoFisher), followed by secondary staining using IRDye800-conjugated anti-mouse secondary. Rho-GTPase immunoprecipitation pull-down was conducted using Glutathione-GST-Rhotekin bait according to manufacturer’s instructions (Thermo, cat# 16116). Bands were quantified using Biorad ImageLab software.

### Enzyme-linked immunosorbent assay

For STMN2 ELISA, 96-well opaque white Maxisorp plates (LifeTechnologies) were coated overnight using Rabbit anti-STMN2 antibody (Abcam, #EPR15286) diluted 1:2000 in carbonate coating buffer. Plates were then washed using PBS-tween and non-specific binding blocked using 1% BSA for 1hr. Samples lysed in RIPA buffer were loaded and incubated at 4°C overnight. Plates were then washed using PBS-tween, and detection antibody (Mouse anti-STMN2, R&D Systems, MAB6930) added diluted 1:1000 in 1% BSA for 1hr at RT. Plates were then washed using PBS-tween, and HRP-conjugated secondary antibody (Novus Biologicals, NB7521) added diluted 1:1000 in 1% BSA for 30 mins at RT. Plates were then washed with PBS-tween, and 100μl SuperSignal ELISA Pico Chemiluminescent substrate added according to manufacturer’s instructions (Thermo). Plates were then read using a BioTek Synergy HTX plate reader at 450nm, and data points normalised to average of control conditions.

### Immunofluorescence and imaging

Cells were plated on 96-well black-walled glass-bottom plates pre-coated with Matrigel (Corning) substrate, prior to 24hr treatment with compounds indicated. Cells were then washed with DPBS with MgCl and CaCl prior to fixation for 15 mins using 3% paraformaldehyde in DPBS. After fixation, cells were permeabilized for 10 mins using 0.25% Triton-X in PBS, washed with DPBS, and blocked using normal serum from the secondary antibody host species. Primary antibodies **(Supplementary Table 8)** were then applied overnight at 4°C, cells were washed using DPBS and then stained for 1hr at room-temperature using fluorescently-conjugated secondary antibodies. Nuclei were stained using either hoescht or DAPI in PBS for 10 mins, followed by washing with PBS and imaging using a Nikon AX confocal microscope. Images were formatted using Fiji (ImageJ). STMN2-mGL expression was quantified in WT *STMN2*-mGreenlantern SH-SY5Y after statin treatment using a custom MATLAB stitching and image analysis pipeline (available upon request from Brian Wainger).

### RNA extraction, complementary DNA synthesis and RT-qPCR

For RNA extraction from cultured cells and neurons, direct lysis was applied using Buffer RLT (RNeasy Mini Kit, Qiagen) and RNA purified according to the manufacturer’s instructions. For cDNA synthesis, 1 μg total RNA was reverse transcribed using the high-capacity reverse transcription kit (Invitrogen) according to the manufacturers’ instructions. RT–PCR reactions were performed using Q5 High-Fidelity DNA polymerase (NEB) in a MasterCycler PCR machine (Eppendorf). RT–PCR products were separated on 2% polyacrylamide gels and then incubated with SYBR gold (Invitrogen) at RT for 1hr for imaging and analysis. Quantitative real-time PCR was carried out in technical triplicates, using iTaq Universal SYBR green (Bio-Rad) in a CFX384 real-time PCR machine. mRNA expression of glyceraldehyde-3-phosphate dehydrogenase (*GAPDH*) was used as an endogenous control gene, as indicated. For RT-qPCR in iPSC-derived motor neurons, pre-designed TaqMan assays (Thermo) for *STMN2* and *EIF4A2* were used. A full list of the expression primers used in this study is included in **Supplementary Table 9**.

### RNA sequencing

Total RNA was extracted from WT SH-SY5Y cells treated with either 0.5% DMSO, 6μM Simvastatin, or 6μM Simvastatin + 30μM mevalonate. Subsequently, 1μg of total RNA was used for mRNA libraries preparation using the TruSeq RNA kit (Illumina). Five biological replicates were sequenced per condition. Sequencing was performed on an Illumina HiSeq 4000 platform with a median of >25M reads per sample. Fifty-base-pair single-end FASTQ files were obtained using the Illumina demultiplexing pipeline. STAR and RSEM were used to align the reads to the human reference sequence HG38 and to calculate the raw counts and transcripts per million values for genes, respectively **(Supplementary Table 4)**. Genes differentially expressed between sample groups were identified using the DESeq2 package in RStudio **(Supplementary Table 5)**. Gene ontology analysis was conducted using ShinyGO^119^ **(Supplementary Table 6)**. Background genes were set as all genes identified through DESeq that produced a padj value. Up/downregulated genes were all genes with a padj <0.05 and a positive or negative log-fold change. Heat-maps were produced using GraphPad Prism. Volcano plots were produced in RStudio using DEseq data with an additional filter of expressed genes as described previously ^120^ and ‘lfcShrink’ ashr function.^121^

### Palmitoylation Detection using Acyl-Biotinyl Exchange (ABE)

Palmitoylated proteins were detected using three steps as adapted from Wan et al:^57^ 1) Blockage of free cysteine thiol groups. 0.1% DMSO and 1μM cerivastatin treated SH-SY5Y cell pellets were solubilized in lysis buffer (50 mM HEPES pH 7.0, 2% SDS (w/v), 1 mM EDTA) containing protease inhibitors and 25 mM N-Ethylmaleimide (NEM) incubated at room temperature with end over end mixing overnight. NEM was removed by acetone precipitation with pellet resuspension in 4SB buffer (4% SDS, 50 mM Tris, 5 mM EDTA, pH 7.4). 2) Removal of thioester linked palmitoyl groups by hydroxylamine (HAM). The resuspended protein sample was split into equal volumes that were incubated with 50 mM Tris (negative control) or 0.7 mM HAM at room temperature for 15 min with end-over-end mixing. 3) Biotinylation of previously palmitoylated sulfhydryl groups. Samples were incubated with sulfhydryl specific HPDP-biotin (0.4 mM, 50 mM Tris-HCl, pH 7.4) for 1 h at room temperature with end over end mixing. Unbound biotin was removed using three sequential acetone precipitations and biotinylated proteins were extracted using NeutrAvidin resin. Bound proteins were eluted, subjected to SDS-PAGE, and Stathmin-2 protein detected by immunoblotting with Rabbit polyclonal anti-STMN2 (Proteintech, #10586-1-AP).

### Dual-gRNA CRISPR inhibition in NGN2 cortical neurons

Dual gRNAs for *MVD* and *GGPS1* **(Supplementary Table 8)** were cloned as a single gBlock fragment into N18-pKLV2-EF1a-BSD-BFP backbone. This is a tRNA dual gRNA system co-expressing BFP and blasticidin resistance. Clones were then integrated into lentivirus and transduced into WTC11^122,123^ iPSC bearing the integrated CRISPRi machinery at day 0 for 48 hours. These cells also featured a stable, doxycycline-inducible *NGN2* expression system used to drive differentiation into cortical neurons^62^. After blasticidin selection, cells were exposed to doxycycline for 11 days using the culture conditions described above prior to harvesting and RNA extraction.

### CRISPR-Cas9 ATF3 knockout line

WT SH-SY5Y cells were cultured as above. The *ATF3* gene was knocked out using a single guide RNA **(Supplementary Table 8)** and WT Cas9 RNP complex transfected into WT SH-SY5Y. The gRNA was designed using CHOPCHOP tool.^124^ After transfection, cells were selected using serial dilution and the target region sequenced in isolated clones. Sequencing revealed the presence of a homozygous single base-pair deletion in exon 1 leading to the formation of a premature stop codon 17bp downstream of the deletion predicted to induce non-sense mediated decay and to prevent translation of the DNA-binding domain **(Extended data Fig. 4a)**. Sequencing after characterization of the line revealed the indel was maintained in >98% reads and therefore not selected out through mitosis **(Extended data Fig. 4b.)**.

### Mice/Animal Care

All animal care and experimental procedures were performed in accordance with animal study proposals approved by the National Institute of Child Health and Human Development Animal Care and Use Committee, animal protocol number ASP 23-003. Atf3^fl/fl^ homozygous mice (Dr. Clifford Woolf lab)^125^ were crossed to the Chat-IRES-Cre::deltaNeo line (Chat^tm1(cre)Lowl^/J); Jax Stock No. 031661). These mice were bred to generate a male breeder with the genotype Atf3^fl/fl^;Chat-iCre+/wt who was then crossed with an Atf3^fl/fl^ female breeder to generate an experimental cohort of roughly 50% Atf3cKO mice (Atf3^fl/fl^;Chat-iCre^+/wt^) and 50% Atf3 control littermate mice (Atf3^fl/fl^;Chat-iCre^wt/wt^) mice. Mice were kept in their parent cages until 3 weeks of age, after which they were weaned into cages of up to five mice of the same sex. After weaning mice were place on regular diet (NIH-07 Rodent Chow) and water was available *ad libidum*. A 12hr light/dark cycle was maintained, with lights on from 6:00 a.m. to 5:59 p.m.

### Sciatic Nerve Crush Surgery

Six or 7-week-old mice were anesthetized at 2.0% isoflurane and maintained between 1.5-2.0% isoflurane for the duration of the surgery. The left hindlimb and surrounding area were shaved to remove fur and cleaned with beta-iodine and 70% ethanol. A single incision was made in the skin midway between the sciatic notch and the knee and muscles were parted to expose the sciatic nerve prior to its distal branch point. The sciatic nerve was crushed just proximal to the branch point for 10 seconds using 2SP laminectomy forceps (F.S.T. Item No. 11223-20). The muscle tissues overlaying the injury were gently placed back together and the skin incision was closed using wound clips (F.S.T. Item No. 12020-00 and 12022-09). Wound clips were kept in place until tissue collection 7 days post injury.

### Tissue Collection

Mice were anesthetized with intraperitoneal injections of 2.1% Avertin, then transcardially perfused with 10mL of 1x PBS followed by 10mL of 4% paraformaldehyde (PFA). Spinal cord was collected and post-fixed for 24 hours in 4% PFA before transfer to 1x PBS + 0.01% sodium azide for storage.

### Immunostaining and RNA-ISH

The lumbar region of the spinal cord was isolated and placed in a 30% sucrose solution overnight before being embedded in O.C.T. compound (Tissue-Tek) and frozen on dry ice. Blocks were then sectioned into 16-μm thick coronal slices onto positively charged slides using a Lecia CM3050 Research Cryostat. For immunostaining, spinal cord sections were rinsed in PBS followed by PBS containing 0.1% Triton X (PBSTx). Sections were then blocked in 5% normal donkey serum in PBSTx for 1-2 hours at room temperature then incubated with primary antibody diluted in 0.5% normal donkey serum in PBSTx overnight at 4°C. Slides were washed in PBSTx, then incubated in AlexaFluor-conjugated secondary antibodies (ThermoFisher) in PBSTx for 2 hours at room temperature, followed by washes in PBSTx and PBS. Finally slides were incubated DAPI (4’,6-diamidino-2-phenylindole); ThermoFisher 62248) diluted in PBS before a final PBS wash. Slides were coverslipped using Prolong Diamond Antifade Mountant (Invitrogen #P36970). Primary Antibodies: anti-STMN2 (Novus Biologicals #NBP1-49461, 1:1000), anti-VAChT (Synaptic Systems #139105, 1:500), anti-ATF3 (Santa Cruz #sc-81189, 1:250). Multiplexed *in situ* hybridization on spinal cord sections was performed according to the manufacturer’s instructions for fixed frozen sections (ACD: 323100, 323120) with the addition of an extended bake time of 60 min. Atf3 (426891), Stmn2 (498391-C2) and Chat (408731-C3) probes were used with C2 and C3 probes diluted 1:500 in C1 probe solution.

### Image Analysis

Slides were imaged using either a Zeiss Axiocam 506 color camera or an Olympus VS200 slide scanner. Images were stitched using either Zeiss Zen software (blue edition) or Olyvia (Olympus). FIJI was used to generate sum intensity images for quantification. The experimenter performing analysis was blind to genotype. For each spinal cord section, each motor neuron (on both contralateral and ipsilateral sides) was outlined based on expression of VAChT to produce individual motor neuron ROIs. The STMN2 intensity of each motor neuron ROI was measured for each section. The intensity of the uninjured motor neurons (contralateral and uninjured ipsilateral) was averaged per section to determine the baseline level of STMN2 expression in motor neurons in each section. The injured motor neurons were anatomically defined as being within the upper motor neuron pool (sciatic) on the injured side, with increased levels of STMN2 and ATF3 expression in WT animals. The STMN2 intensities of the injured motor neurons were each individually divided by the average baseline STMN2 intensity for that section to generate a normalized value for each injured motor neuron. 10 sections with injured motor neurons were quantified per animal per genotype. Statistics are reported on the average of values per animal.

### Statistics

Statistical tests were performed using GraphPad Prism. Student’s *t*-tests (two-tailed or one-tailed), one-way or two-way ANOVA were used as indicated. No statistical methods were used to pre-determine sample sizes but our sample sizes are similar to those reported in previous publications. Experiments were not randomized. Data distribution was tested for normality using D’Agostino and Pearson method when obvious outliers were observed but was otherwise assumed to be normal. Data collection and analysis were not performed blind to the conditions of the experiments. Levels for quoted confidence intervals are set at 95% and *p*-values in the figures are indicated as follows: * *p* < 0.05, ** *p* < 0.01, *** *p* < 0.001. Error bars = standard deviation unless otherwise stated.

## Acknowledgements

We are grateful to all the individuals who donated their blood or fibroblasts for the generation of iPSCs used in this study. We thank all members of the Lagier-Tourenne, Wainger, Albers, Cleveland, Lutz and Ward labs for helpful discussions and support. We are grateful for the support of the MGB Innovation team, in particular David Silva and Tom Backus. Screening was conducted at ICCB-Longwood, Harvard Medical School with the assistance of Jennifer Smith, Richard Siu, Jennifer Splaine, and David Wrobel. Stathmin-2 ELISA protocol kindly provided by Dr. Joseph Klim from Dr. Eggan’s lab. We thank Dr. Tracy Cole and Dr. Benjamin Moore for their help throughout.

## Author contributions

Conceptualization: MN, CLP, CLT

Methodology: MN, SA, ISN, MC, JW, MH, AH, SML, XJ, NR, AAAQ, AB, MB, MP, CA, JH, TC, BM, ZM, BPK, MW, DWC, RT, CML, RH, DK, MW, CRRA, BW, CLP, CLT

Investigation: MN, SA, ISN, MC, JW, MH, SY, PL, HB, BW, CZL, SML, XJ, NR, AAAQ, CJFR, AR, GG, CA, JH, HW

Visualization: MN, SA, ISN, JW, PL, BW, BM, TC

Funding acquisition: MN, DWC, CML, RH, DK, MW, CLT

Supervision: BPK, DWC, RT, CLL, MW, CRRA, BW, CLP, CLT

Writing – original draft: MN, CLT

Writing – reviewing & editing: All authors

## Data and materials availability

Generated lines are available upon material transfer agreement with Massachusetts General Hospital. Code used in this study can be found at: https://github.com/waingerlab/Nolan_etal.

---

**Extended data Fig 1.**
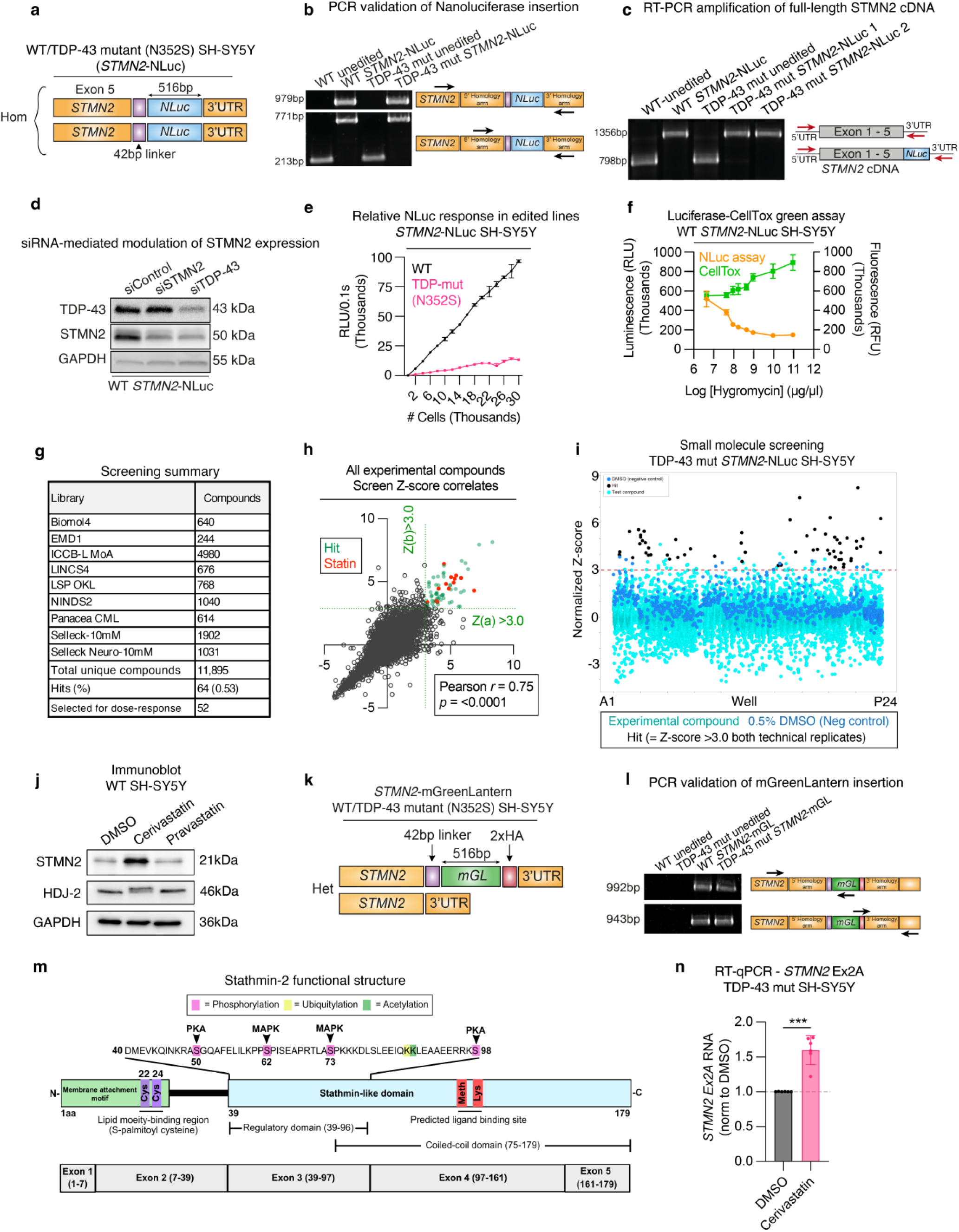
**(A)** Schematic of *STMN2*-NLuc models. WT and TDP-43 mutant (N352S) SH-SY5Y cells were CRISPR-Cas9 edited to express Nanoluciferase at the C-terminus of both endogenous *STMN2* alleles. **(B)** PCR of edited lines using primers outside and within the repair template homology arms confirms homozygosity of the insertion. **(C)** RT-PCR using primers spanning the entire *STMN2* cDNA demonstrate no unusual splice variants after NLuc CRISPR-Cas9 insertion. **(D)** Immunoblot of WT *STMN2*-NLuc line demonstrates that the sensitivity to STMN2/TDP-43 siRNA-mediated disruption is maintained after editing. **(E)** WT STMN2-NLuc and TDP-43 mutant (N352S) lines demonstrate linear luciferase response up to 20k plated cells proportional to TDP-43 status. **(F)** Luciferase expression in WT *STMN2*-NLuc line is inversely proportional to cell death (CellTox Green cytotoxicity assay, Promega) after treatment with high-doses of hygromycin. **(G)** Small molecules libraries used for pharmacological screen in *STMN2*-NLuc TDP-43 mutant (N352S) SH-SY5Y cells. **(H-I)** Pearson correlation of replicates **(H)** and Z-score dot plot **(I)** of experimental compounds across the screen. Points in both graphs represent average of technical duplicates. **(J)** Electromobility shift assay (EMSA) using HDJ2 antibody demonstrates the inhibition of HDJ2 farnesylation following cerivastatin, but not pravastatin, treatment which can be visualised as an accumulation of the unprenylated protein and upward shift on immunoblot. **(K)** Schematic for WT and TDP-43 *STMN2*-mGreenLantern lines. **(L)** PCR using primers within and outside the repair template homology arms confirm correct insertion of mGreenLantern at the C-terminal of STMN2 after CRISPR-Cas9 mediated editing. **(M)** Exons and functional domains schematic of STMN2 transcript and protein showing known post-translational modifications. **(N)** RT-qPCR for *STMN2* cryptic exon (Ex2A) expression in TDP-43 mutant (N352S) SH-SY5Y treated with cerivastatin. Points represent mean of technical replicates from independent experiments. Unpaired t-test.

**Extended data Fig 2.**
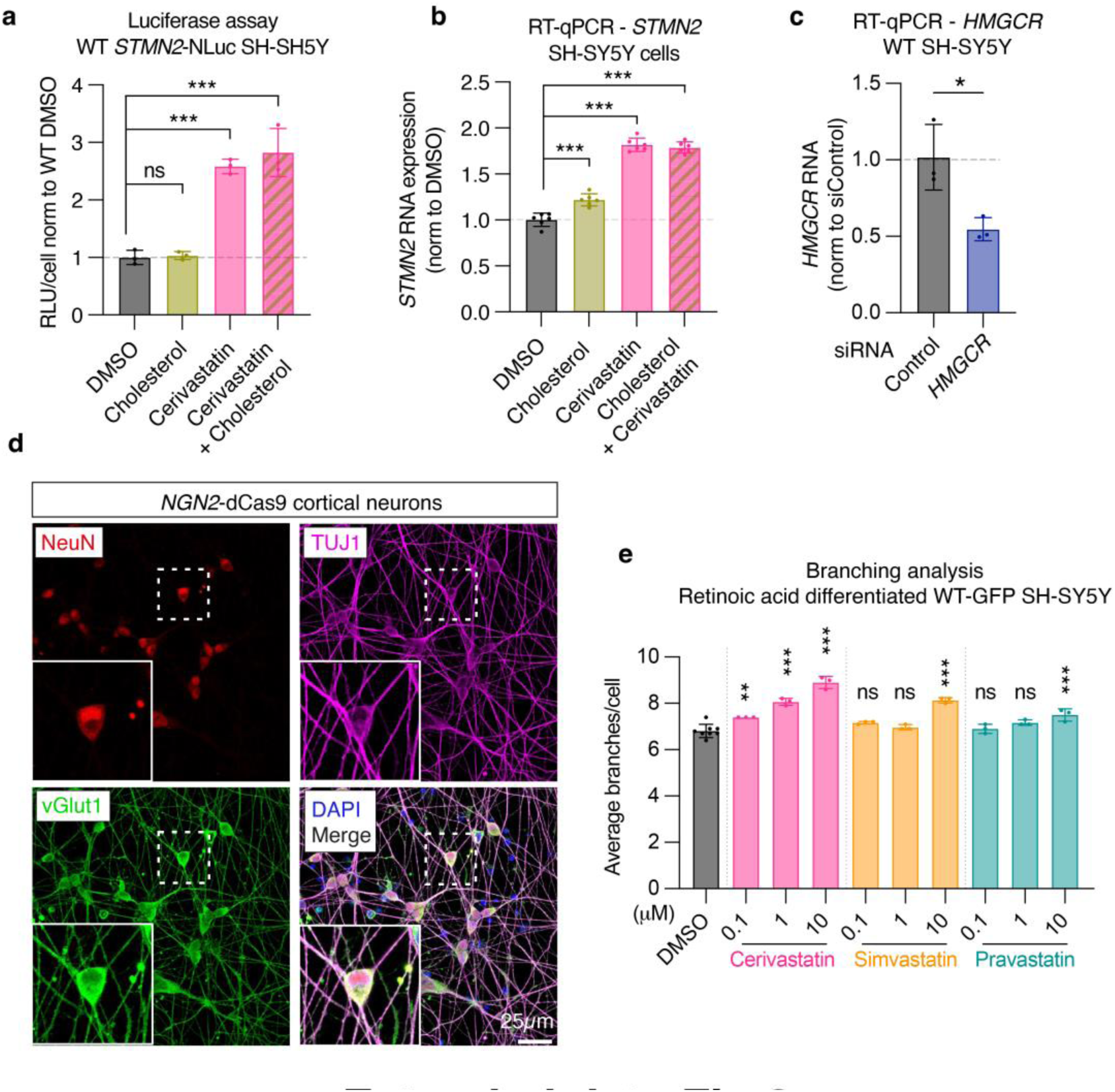
**(A-B)** Luciferase assay in WT STMN2-NLuc cells **(A)** and RT-qPCR in WT SH-SY5Y **(B)** treated with DMSO, 5mg/ml cholesterol or 1mM cerivastatin alone or co-treated with cerivastatin and cholesterol. Points represent mean of technical replicates from three independent experiments. One-way ANOVA with Tukey’s multiple comparisons. **(C)** RT-qPCR quantifying siRNA-mediated knockdown of *HMGCR* in WT SH-SY5Y cells compared to control siRNA. Points represent mean of technical replicates from three independent biological replicates. Unpaired t-test. **(D)** Immunofluorescent staining 11 days post-doxycycline induction of *NGN2*-dCas9 cortical neurons using antibodies targeting NeuN (red), vGlut1 (green), and TUJ1 (purple). Scale bar = 25mm. **(E)** Branching analysis conducted on WT-GFP, retinoic acid differentiated SH-SY5Y after treatment with indicated compounds. Comparisons represented to DMSO control. Points represent average of technical replicates from independent experiments. One-way ANOVA with Dunnett’s multiple comparisons.

**Extended data Fig 3.**
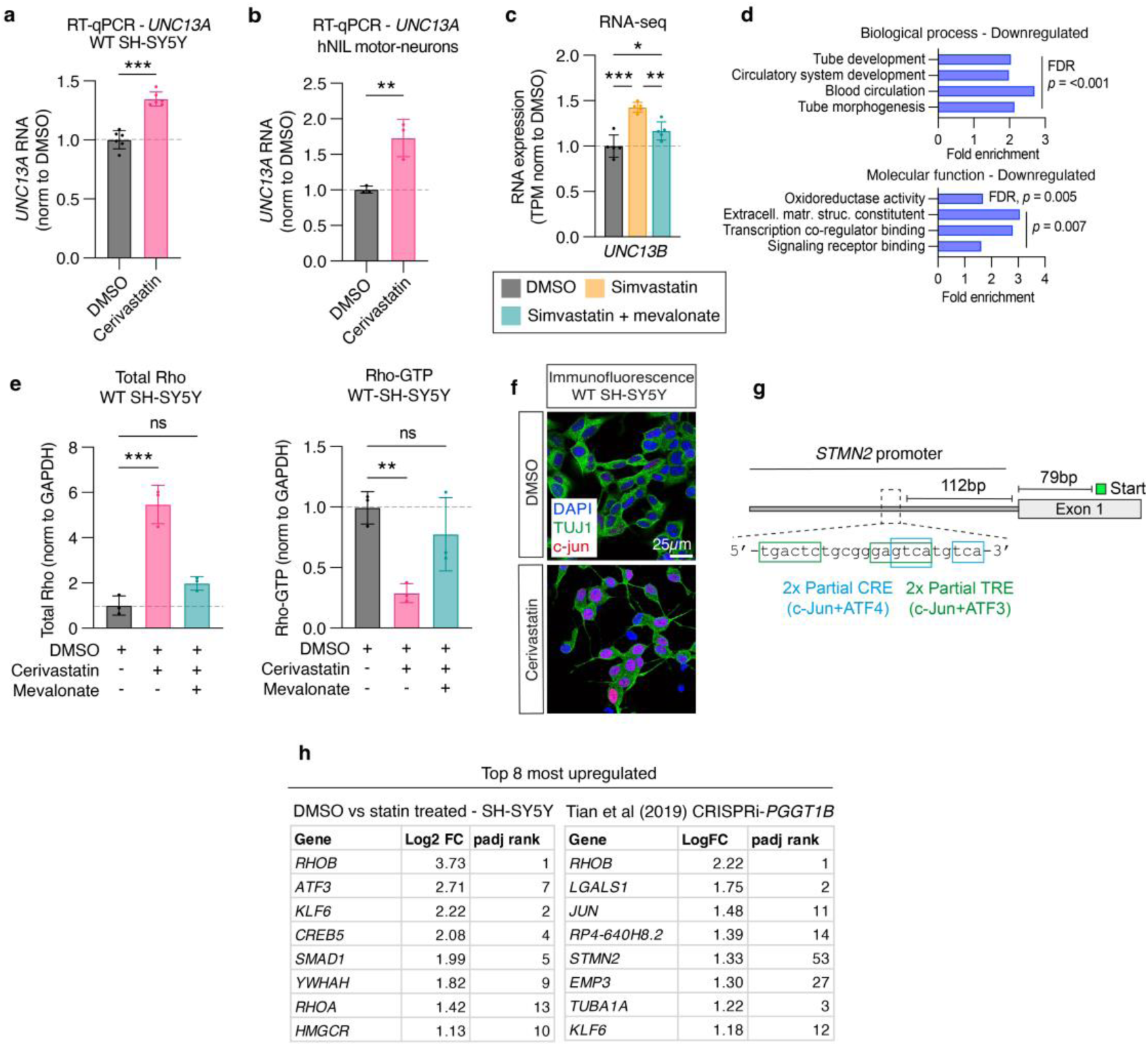
**(A-B)** RT-qPCR for *UNC13A* mRNA in WT SH-SY5Y **(A)** and hNIL motor neurons **(B)** treated with 1mM or 2mM cerivastatin, respectively. Points represent mean of technical replicates from three independent experiments. Unpaired t-tests. **(C)** Levels of *UNC13B* transcripts detected by RNAseq in WT SH-SY5Y treated with DMSO, simvastatin or simvastatin + mevalonate. TPM normalized to DMSO average. Asterisks indicate adjusted p-values obtained using DESeq2. **(D)** Top gene ontology terms identified as enriched using ShinyGO in downregulated genes upon simvastatin treatment. **(E)** Quantification of Total Rho and pull-down Rho-GTP from Fig 4G. WT SH-SY5Y treated with indicated compounds for 24hrs. Points represent replicates from independent experiments. Both graphs one-way ANOVA with Dunnett’s multiple comparisons. **(F)** Immunofluorescent staining of WT SH-SY5Y with c-jun (red) and TUJ1 (green) antibodies following DMSO or 1mM cerivastatin treatment. **(G)** Schematic denoting predicted AP-1 binding sites in the proximal human *STMN2* promoter. **(H)** Top genes (ranked by Log2 FC) upregulated by both statin treatment and CRISPRi-*PGGT1B. RHOB, JUN, STMN2,* and *KLF6* were among the most upregulated in both datasets, with significant upregulations of *RHOB* and *KLF6* maintained in mature iPSC-neurons following *PGGT1B* knockdown^63^.

**Extended data Fig 4.**
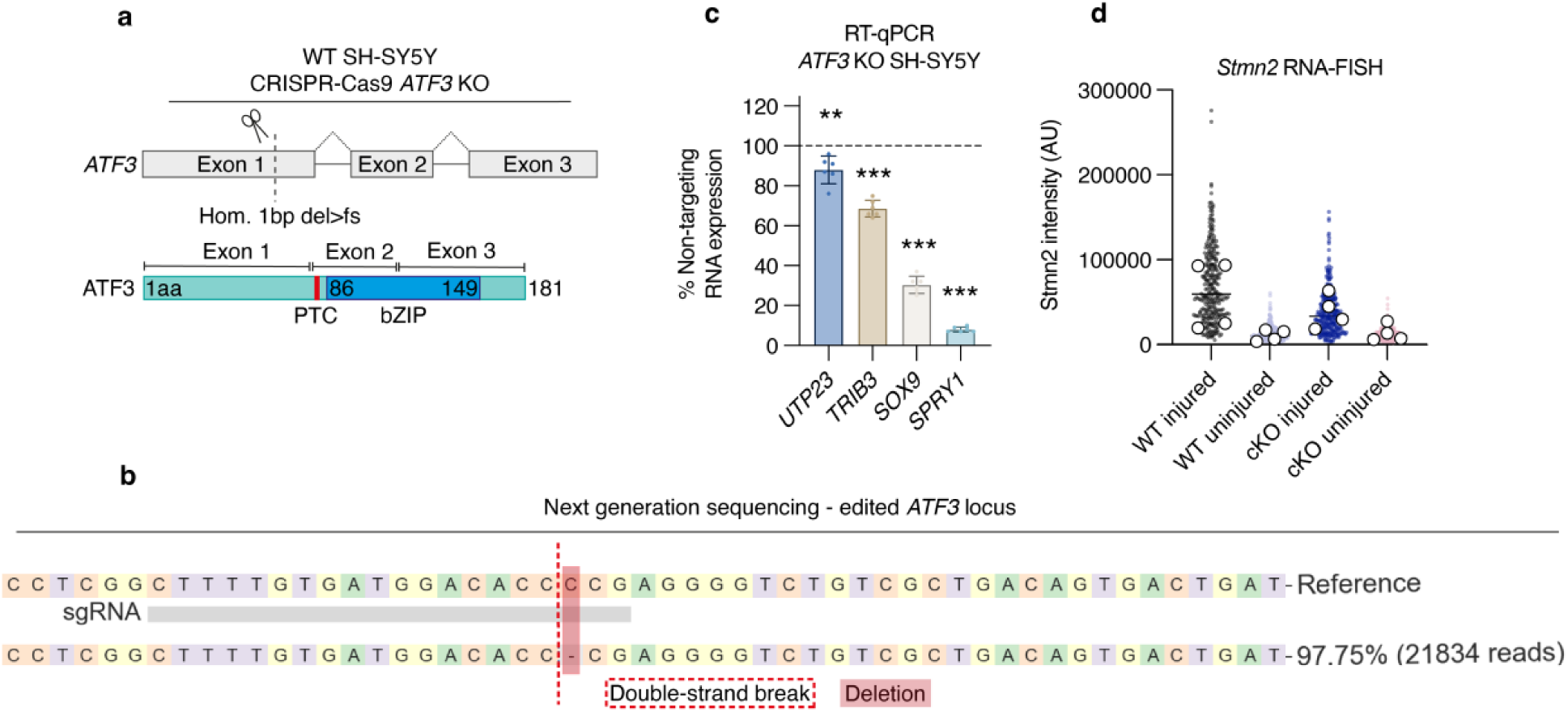
**(A)** Cas9 and a gRNA targeting *ATF3* Exon 1 were used to generate a clone of WT SH-SY5Y with a deletion of 1bp, resulting in a frameshift and premature stop codon upstream of the bZIP DNA-binding domain of ATF3 protein. **(B)** Next-generation sequencing of the targeted locus confirmed the deletion in ∼98% of reads after clonal selection. Sequencing was repeated after characterization of the line to re-confirm the indel. **(C)** RT-qPCR of known *ATF3* targets reveals a downregulation of their RNA in *ATF3* KO line. RNA levels are normalized to values in WT SH-SY5Y generated using a non-targeting sgRNA as control. Points represent mean of technical replicates from independent experiments. Asterisks indicate unpaired t-test comparisons to control cells. **(D)** Quantification of *Stmn2* RNA-FISH in uninjured and injured motor neuron pools from both control and Atf3 cKO mice. Small points represent individual motor neurons; large points represent average for each animal. Bar represents median value.

## References

1 Neumann, M. et al. Ubiquitinated TDP-43 in frontotemporal lobar degeneration and amyotrophic lateral sclerosis. Science 314, 130–133 (2006). 10.1126/science.1134108

2 Arai, T. et al. TDP-43 is a component of ubiquitin-positive tau-negative inclusions in frontotemporal lobar degeneration and amyotrophic lateral sclerosis. Biochem Biophys Res Commun 351, 602–611 (2006). 10.1016/j.bbrc.2006.10.093

3 Nolan, M. et al. Quantitative patterns of motor cortex proteinopathy across ALS genotypes. Acta Neuropathol Commun 8, 98 (2020). 10.1186/s40478-020-00961-2

4 Nelson, P. T. et al. Limbic-predominant age-related TDP-43 encephalopathy (LATE): consensus working group report. Brain 142, 1503–1527 (2019). 10.1093/brain/awz099

5 Josephs, K. A. et al. TDP-43 is a key player in the clinical features associated with Alzheimer’s disease. Acta Neuropathol 127, 811–824 (2014). 10.1007/s00401-014-1269-z

6 Sun, M. et al. Cryptic exon incorporation occurs in Alzheimer’s brain lacking TDP-43 inclusion but exhibiting nuclear clearance of TDP-43. Acta Neuropathol 133, 923–931 (2017). 10.1007/s00401-017-1701-2

7 Vatsavayai, S. C. et al. Timing and significance of pathological features in C9orf72 expansion-associated frontotemporal dementia. Brain 139, 3202–3216 (2016). 10.1093/brain/aww250

8 Nana, A. L. et al. Neurons selectively targeted in frontotemporal dementia reveal early stage TDP-43 pathobiology. Acta Neuropathol 137, 27–46 (2019). 10.1007/s00401-018-1942-8

9 Lagier-Tourenne, C. et al. Divergent roles of ALS-linked proteins FUS/TLS and TDP-43 intersect in processing long pre-mRNAs. Nat Neurosci 15, 1488–1497 (2012). 10.1038/nn.3230

10 Tollervey, J. R. et al. Characterizing the RNA targets and position-dependent splicing regulation by TDP-43. Nat Neurosci 14, 452–458 (2011). 10.1038/nn.2778

11 Polymenidou, M. et al. Long pre-mRNA depletion and RNA missplicing contribute to neuronal vulnerability from loss of TDP-43. Nat Neurosci 14, 459–468 (2011). 10.1038/nn.2779

12 Freibaum, B. D., Chitta, R. K., High, A. A. & Taylor, J. P. Global analysis of TDP-43 interacting proteins reveals strong association with RNA splicing and translation machinery. J Proteome Res 9, 1104–1120 (2010). 10.1021/pr901076y

13 Lagier-Tourenne, C. & Cleveland, D. W. Rethinking ALS: the FUS about TDP-43. Cell 136, 1001–1004 (2009). 10.1016/j.cell.2009.03.006

14 Melamed, Z. et al. Premature polyadenylation-mediated loss of stathmin-2 is a hallmark of TDP-43-dependent neurodegeneration. Nat Neurosci 22, 180–190 (2019). 10.1038/s41593-018-0293-z

15 Klim, J. R. et al. ALS-implicated protein TDP-43 sustains levels of STMN2, a mediator of motor neuron growth and repair. Nat Neurosci 22, 167–179 (2019). 10.1038/s41593-018-0300-4

16 Prudencio, M. et al. Truncated stathmin-2 is a marker of TDP-43 pathology in frontotemporal dementia. J Clin Invest 130, 6080–6092 (2020). 10.1172/JCI139741

17 Pickles, S. et al. Evidence of cerebellar TDP-43 loss of function in FTLD-TDP. Acta Neuropathol Commun 10, 107 (2022). 10.1186/s40478-022-01408-6

18 Agra Almeida Quadros, A. R., et al. Cryptic splicing of stathmin-2 and UNC13A mRNAs is a pathological hallmark of TDP-43-associated Alzheimer’s disease. Acta Neuropathol 147, 9 (2024). 10.1007/s00401-023-02655-0

19 Chung, M. et al. Cryptic exon inclusion is a molecular signature of LATE-NC in aging brains. Acta Neuropathol 147, 29 (2024). 10.1007/s00401-023-02671-0

20 Estades Ayuso, V., et al. TDP-43-regulated cryptic RNAs accumulate in Alzheimer’s disease brains. Mol Neurodegener 18, 57 (2023). 10.1186/s13024-023-00646-z

21 Holmfeldt, P., Brannstrom, K., Stenmark, S. & Gullberg, M. Deciphering the cellular functions of the Op18/Stathmin family of microtubule-regulators by plasma membrane-targeted localization. Mol Biol Cell 14, 3716–3729 (2003). 10.1091/mbc.e03-03-0126

22 Charbaut, E. et al. Stathmin family proteins display specific molecular and tubulin binding properties. J Biol Chem 276, 16146–16154 (2001). 10.1074/jbc.M010637200

23 Curmi, P. A. et al. Stathmin and its phosphoprotein family: general properties, biochemical and functional interaction with tubulin. Cell Struct Funct 24, 345–357 (1999). 10.1247/csf.24.345

24 Grenningloh, G., Soehrman, S., Bondallaz, P., Ruchti, E. & Cadas, H. Role of the microtubule destabilizing proteins SCG10 and stathmin in neuronal growth. J Neurobiol 58, 60–69 (2004). 10.1002/neu.10279

25 Morii, H., Shiraishi-Yamaguchi, Y. & Mori, N. SCG10, a microtubule destabilizing factor, stimulates the neurite outgrowth by modulating microtubule dynamics in rat hippocampal primary cultured neurons. J Neurobiol 66, 1101–1114 (2006). 10.1002/neu.20295

26 Baughn, M. W. et al. Mechanism of STMN2 cryptic splice-polyadenylation and its correction for TDP-43 proteinopathies. Science 379, 1140–1149 (2023). 10.1126/science.abq5622

27 Lopez-Erauskin, J. et al. Stathmin-2 loss leads to neurofilament-dependent axonal collapse driving motor and sensory denervation. Nat Neurosci 27, 34–47 (2024). 10.1038/s41593-023-01496-0

28 Guerra San Juan, I., et al. Loss of mouse Stmn2 function causes motor neuropathy. Neuron 110, 1671–1688 e1676 (2022). 10.1016/j.neuron.2022.02.011

29 Krus, K. L. et al. Loss of Stathmin-2, a hallmark of TDP-43-associated ALS, causes motor neuropathy. Cell Rep 39, 111001 (2022). 10.1016/j.celrep.2022.111001

30 Tobert, J. A. Lovastatin and beyond: the history of the HMG-CoA reductase inhibitors. Nat Rev Drug Discov 2, 517–526 (2003). 10.1038/nrd1112

31 Istvan, E. S. & Deisenhofer, J. Structural mechanism for statin inhibition of HMG-CoA reductase. Science 292, 1160–1164 (2001). 10.1126/science.1059344

32 Fracassi, A. et al. Statins and the Brain: More than Lipid Lowering Agents? Curr Neuropharmacol 17, 59–83 (2019). 10.2174/1570159X15666170703101816

33 Greenwood, J., Steinman, L. & Zamvil, S. S. Statin therapy and autoimmune disease: from protein prenylation to immunomodulation. Nat Rev Immunol 6, 358–370 (2006). 10.1038/nri1839

34 Liao, J. K. Isoprenoids as mediators of the biological effects of statins. J Clin Invest 110, 285–288 (2002). 10.1172/JCI16421

35 Zhang, F. L. & Casey, P. J. Protein prenylation: molecular mechanisms and functional consequences. Annu Rev Biochem 65, 241–269 (1996). 10.1146/annurev.bi.65.070196.001325

36 Hori, Y. et al. Post-translational modifications of the C-terminal region of the rho protein are important for its interaction with membranes and the stimulatory and inhibitory GDP/GTP exchange proteins. Oncogene 6, 515–522 (1991).

37 Li, X. et al. Inhibition of protein geranylgeranylation and RhoA/RhoA kinase pathway induces apoptosis in human endothelial cells. J Biol Chem 277, 15309–15316 (2002). 10.1074/jbc.M201253200

38 Marinissen, M. J., Chiariello, M. & Gutkind, J. S. Regulation of gene expression by the small GTPase Rho through the ERK6 (p38 gamma) MAP kinase pathway. Genes Dev 15, 535–553 (2001). 10.1101/gad.855801

39 Marinissen, M. J. et al. The small GTP-binding protein RhoA regulates c-jun by a ROCK-JNK signaling axis. Mol Cell 14, 29–41 (2004). 10.1016/s1097-2765(04)00153-4

40 Dupraz, S. et al. RhoA Controls Axon Extension Independent of Specification in the Developing Brain. Curr Biol 29, 3874–3886 e3879 (2019). 10.1016/j.cub.2019.09.040

41 Stern, S. et al. RhoA drives actin compaction to restrict axon regeneration and astrocyte reactivity after CNS injury. Neuron 109, 3436–3455 e3439 (2021). 10.1016/j.neuron.2021.08.014

42 Li, H. et al. Protein Prenylation Constitutes an Endogenous Brake on Axonal Growth. Cell Rep 16, 545–558 (2016). 10.1016/j.celrep.2016.06.013

43 Morimoto, S. et al. Phase 1/2a clinical trial in ALS with ropinirole, a drug candidate identified by iPSC drug discovery. Cell Stem Cell 30, 766–780 e769 (2023). 10.1016/j.stem.2023.04.017

44 Whitlon, D. S. et al. Novel High Content Screen Detects Compounds That Promote Neurite Regeneration from Cochlear Spiral Ganglion Neurons. Sci Rep 5, 15960 (2015). 10.1038/srep15960

45 Shabanzadeh, A. P. et al. Cholesterol synthesis inhibition promotes axonal regeneration in the injured central nervous system. Neurobiol Dis 150, 105259 (2021). 10.1016/j.nbd.2021.105259

46 Jin, Y. et al. Atorvastatin enhances neurite outgrowth in cortical neurons in vitro via up-regulating the Akt/mTOR and Akt/GSK-3beta signaling pathways. Acta Pharmacol Sin 33, 861–872 (2012). 10.1038/aps.2012.59

47 Iwamoto, K., Yoshii, Y. & Ikeda, K. Atorvastatin treatment attenuates motor neuron degeneration in wobbler mice. Amyotroph Lateral Scler 10, 405–409 (2009). 10.3109/17482960902870993

48 McTaggart, F. et al. Preclinical and clinical pharmacology of Rosuvastatin, a new 3-hydroxy-3-methylglutaryl coenzyme A reductase inhibitor. Am J Cardiol 87, 28B–32B (2001). 10.1016/s0002-9149(01)01454-0

49 Murphy, C., Deplazes, E., Cranfield, C. G. & Garcia, A. The Role of Structure and Biophysical Properties in the Pleiotropic Effects of Statins. Int J Mol Sci 21 (2020). 10.3390/ijms21228745

50 Fong, C. W. Statins in therapy: understanding their hydrophilicity, lipophilicity, binding to 3-hydroxy-3-methylglutaryl-CoA reductase, ability to cross the blood brain barrier and metabolic stability based on electrostatic molecular orbital studies. Eur J Med Chem 85, 661–674 (2014). 10.1016/j.ejmech.2014.08.037

51 Hsiang, B. et al. A novel human hepatic organic anion transporting polypeptide (OATP2). Identification of a liver-specific human organic anion transporting polypeptide and identification of rat and human hydroxymethylglutaryl-CoA reductase inhibitor transporters. J Biol Chem 274, 37161–37168 (1999). 10.1074/jbc.274.52.37161

52 Berndt, N. & Sebti, S. M. Measurement of protein farnesylation and geranylgeranylation in vitro, in cultured cells and in biopsies, and the effects of prenyl transferase inhibitors. Nat Protoc 6, 1775–1791 (2011). 10.1038/nprot.2011.387

53 Chauvin, S., Poulain, F. E., Ozon, S. & Sobel, A. Palmitoylation of stathmin family proteins domain A controls Golgi versus mitochondrial subcellular targeting. Biol Cell 100, 577–589 (2008). 10.1042/BC20070119

54 Lutjens, R. et al. Localization and targeting of SCG10 to the trans-Golgi apparatus and growth cone vesicles. Eur J Neurosci 12, 2224–2234 (2000). 10.1046/j.1460-9568.2000.00112.x

55 Charbaut, E., Chauvin, S., Enslen, H., Zamaroczy, S. & Sobel, A. Two separate motifs cooperate to target stathmin-related proteins to the Golgi complex. J Cell Sci 118, 2313–2323 (2005). 10.1242/jcs.02349

56 Hovde, M. J. et al. Sodium hydrogen exchanger (NHE1) palmitoylation and potential functional regulation. Life Sci 288, 120142 (2022). 10.1016/j.lfs.2021.120142

57 Wan, J., Roth, A. F., Bailey, A. O. & Davis, N. G. Palmitoylated proteins: purification and identification. Nat Protoc 2, 1573–1584 (2007). 10.1038/nprot.2007.225

58 Liao, J. K. & Laufs, U. Pleiotropic effects of statins. Annu Rev Pharmacol Toxicol 45, 89–118 (2005). 10.1146/annurev.pharmtox.45.120403.095748

59 Davignon, J. Beneficial cardiovascular pleiotropic effects of statins. Circulation 109, III39–43 (2004). 10.1161/01.CIR.0000131517.20177.5a

60 Suzumura, K., Yasuhara, M., Tanaka, K. & Suzuki, T. Protective effect of fluvastatin sodium (XU-62-320), a 3-hydroxy-3-methylglutaryl coenzyme A (HMG-CoA) reductase inhibitor, on oxidative modification of human low-density lipoprotein in vitro. Biochem Pharmacol 57, 697–703 (1999). 10.1016/s0006-2952(98)00341-4

61 Ridker, P. M. et al. Inflammation, pravastatin, and the risk of coronary events after myocardial infarction in patients with average cholesterol levels. Cholesterol and Recurrent Events (CARE) Investigators. Circulation 98, 839–844 (1998). 10.1161/01.cir.98.9.839

62 Fernandopulle, M. S. et al. Transcription Factor-Mediated Differentiation of Human iPSCs into Neurons. Curr Protoc Cell Biol 79, e51 (2018). 10.1002/cpcb.51

63 Tian, R. et al. CRISPR Interference-Based Platform for Multimodal Genetic Screens in Human iPSC-Derived Neurons. Neuron 104, 239–255 e212 (2019). 10.1016/j.neuron.2019.07.014

64 Zandl-Lang, M. et al. Regulatory effects of simvastatin and apoJ on APP processing and amyloid-beta clearance in blood-brain barrier endothelial cells. Biochim Biophys Acta Mol Cell Biol Lipids 1863, 40–60 (2018). 10.1016/j.bbalip.2017.09.008

65 Robin, N. C. et al. Simvastatin promotes adult hippocampal neurogenesis by enhancing Wnt/beta-catenin signaling. Stem Cell Reports 2, 9–17 (2014). 10.1016/j.stemcr.2013.11.002

66 Ma, X. R. et al. TDP-43 represses cryptic exon inclusion in the FTD-ALS gene UNC13A. Nature 603, 124–130 (2022). 10.1038/s41586-022-04424-7

67 Brown, A. L. et al. TDP-43 loss and ALS-risk SNPs drive mis-splicing and depletion of UNC13A. Nature 603, 131–137 (2022). 10.1038/s41586-022-04436-3

68 Mahar, M. & Cavalli, V. Intrinsic mechanisms of neuronal axon regeneration. Nat Rev Neurosci 19, 323–337 (2018). 10.1038/s41583-018-0001-8

69 Nguyen, M. Q., Le Pichon, C. E. & Ryba, N. Stereotyped transcriptomic transformation of somatosensory neurons in response to injury. Elife 8 (2019). 10.7554/eLife.49679

70 Seijffers, R. et al. ATF3 expression improves motor function in the ALS mouse model by promoting motor neuron survival and retaining muscle innervation. Proc Natl Acad Sci U S A 111, 1622–1627 (2014). 10.1073/pnas.1314826111

71 Shin, J. E., Geisler, S. & DiAntonio, A. Dynamic regulation of SCG10 in regenerating axons after injury. Exp Neurol 252, 1–11 (2014). 10.1016/j.expneurol.2013.11.007

72 Kim, H. W. et al. Long-term tactile hypersensitivity after nerve crush injury in mice is characterized by the persistence of intact sensory axons. Pain 164, 2327–2342 (2023). 10.1097/j.pain.0000000000002937

73 Dubovy, P., Klusakova, I., Hradilova-Svizenska, I. & Joukal, M. Expression of Regeneration-Associated Proteins in Primary Sensory Neurons and Regenerating Axons After Nerve Injury-An Overview. Anat Rec (Hoboken) 301, 1618–1627 (2018). 10.1002/ar.23843

74 Eferl, R. & Wagner, E. F. AP-1: a double-edged sword in tumorigenesis. Nat Rev Cancer 3, 859–868 (2003). 10.1038/nrc1209

75 Wlaschin, J. J. et al. Promoting regeneration while blocking cell death preserves motor neuron function in a model of ALS. Brain 146, 2016–2028 (2023). 10.1093/brain/awac415

76 Choi, S. H. et al. A three-dimensional human neural cell culture model of Alzheimer’s disease. Nature 515, 274–278 (2014). 10.1038/nature13800

77 Zhao, J., Li, X., Guo, M., Yu, J. & Yan, C. The common stress responsive transcription factor ATF3 binds genomic sites enriched with p300 and H3K27ac for transcriptional regulation. BMC Genomics 17, 335 (2016). 10.1186/s12864-016-2664-8

78 Lu, X. et al. Identification of ATF3 as a novel protective signature of quiescent colorectal tumor cells. Cell Death Dis 14, 676 (2023). 10.1038/s41419-023-06204-1

79 Wlaschin, J. J. et al. Dual leucine zipper kinase is required for mechanical allodynia and microgliosis after nerve injury. Elife 7 (2018). 10.7554/eLife.33910

80 Shin, J. E. et al. SCG10 is a JNK target in the axonal degeneration pathway. Proc Natl Acad Sci U S A 109, E3696–3705 (2012). 10.1073/pnas.1216204109

81 Chauvin, S. & Sobel, A. Neuronal stathmins: a family of phosphoproteins cooperating for neuronal development, plasticity and regeneration. Prog Neurobiol 126, 1–18 (2015). 10.1016/j.pneurobio.2014.09.002

82 Gavet, O., El Messari, S., Ozon, S. & Sobel, A. Regulation and subcellular localization of the microtubule-destabilizing stathmin family phosphoproteins in cortical neurons. J Neurosci Res 68, 535–550 (2002). 10.1002/jnr.10234

83 Stein, R., Orit, S. & Anderson, D. J. The induction of a neural-specific gene, SCG10, by nerve growth factor in PC12 cells is transcriptional, protein synthesis dependent, and glucocorticoid inhibitable. Dev Biol 127, 316–325 (1988). 10.1016/0012-1606(88)90318-1

84 Abdolmaleki, A., Zahri, S. & Bayrami, A. Rosuvastatin enhanced functional recovery after sciatic nerve injury in the rat. Eur J Pharmacol 882, 173260 (2020). 10.1016/j.ejphar.2020.173260

85 Pan, H. C. et al. Neuroprotective effect of atorvastatin in an experimental model of nerve crush injury. Neurosurgery 67, 376–388; discussion 388-379 (2010). 10.1227/01.NEU.0000371729.47895.A0

86 Kreple, C. J. et al. Protective Effects of Lovastatin in a Population-Based ALS Study and Mouse Model. Ann Neurol (2023). 10.1002/ana.26600

87 Colman, E. et al. An evaluation of a data mining signal for amyotrophic lateral sclerosis and statins detected in FDA’s spontaneous adverse event reporting system. Pharmacoepidemiol Drug Saf 17, 1068–1076 (2008). 10.1002/pds.1643

88 Golomb, B. A., Verden, A., Messner, A. K., Koslik, H. J. & Hoffman, K. B. Amyotrophic Lateral Sclerosis Associated with Statin Use: A Disproportionality Analysis of the FDA’s Adverse Event Reporting System. Drug Saf 41, 403–413 (2018). 10.1007/s40264-017-0620-4

89 Torrandell-Haro, G. et al. Statin therapy and risk of Alzheimer’s and age-related neurodegenerative diseases. Alzheimers Dement (N Y) 6, e12108 (2020). 10.1002/trc2.12108

90 Jeong, S. M. et al. Association between statin use and Alzheimer’s disease with dose response relationship. Sci Rep 11, 15280 (2021). 10.1038/s41598-021-94803-3

91 Poly, T. N. et al. Association between Use of Statin and Risk of Dementia: A Meta-Analysis of Observational Studies. Neuroepidemiology 54, 214–226 (2020). 10.1159/000503105

92 Ren, Q. W. et al. Statins and risks of dementia among patients with heart failure: a population-based retrospective cohort study in Hong Kong. Lancet Reg Health West Pac 44, 101006 (2024). 10.1016/j.lanwpc.2023.101006

93 Zhang, X., Wen, J. & Zhang, Z. Statins use and risk of dementia: A dose-response meta analysis. Medicine (Baltimore) 97, e11304 (2018). 10.1097/MD.0000000000011304

94 Olmastroni, E. et al. Statin use and risk of dementia or Alzheimer’s disease: a systematic review and meta-analysis of observational studies. Eur J Prev Cardiol 29, 804–814 (2022). 10.1093/eurjpc/zwab208

95 Pfeiffer, R. M. et al. Identifying potential targets for prevention and treatment of amyotrophic lateral sclerosis based on a screen of medicare prescription drugs. Amyotroph Lateral Scler Frontotemporal Degener 21, 235–245 (2020). 10.1080/21678421.2019.1682613

96 Weisskopf, M. G., Levy, J., Dickerson, A. S., Paganoni, S. & Leventer-Roberts, M. Statin Medications and Amyotrophic Lateral Sclerosis Incidence and Mortality. Am J Epidemiol 191, 1248–1257 (2022). 10.1093/aje/kwac054

97 Freedman, D. M., Kuncl, R. W., Cahoon, E. K., Rivera, D. R. & Pfeiffer, R. M. Relationship of statins and other cholesterol-lowering medications and risk of amyotrophic lateral sclerosis in the US elderly. Amyotroph Lateral Scler Frontotemporal Degener 19, 538–546 (2018). 10.1080/21678421.2018.1511731

98 Jeong, A. et al. Protein farnesylation is upregulated in Alzheimer’s human brains and neuron-specific suppression of farnesyltransferase mitigates pathogenic processes in Alzheimer’s model mice. Acta Neuropathol Commun 9, 129 (2021). 10.1186/s40478-021-01231-5

99 Eckert, G. P. et al. Regulation of the brain isoprenoids farnesyl- and geranylgeranylpyrophosphate is altered in male Alzheimer patients. Neurobiol Dis 35, 251–257 (2009). 10.1016/j.nbd.2009.05.005

100 Jeong, A., Suazo, K. F., Wood, W. G., Distefano, M. D. & Li, L. Isoprenoids and protein prenylation: implications in the pathogenesis and therapeutic intervention of Alzheimer’s disease. Crit Rev Biochem Mol Biol 53, 279–310 (2018). 10.1080/10409238.2018.1458070

101 Jeong, A. et al. In Vivo Prenylomic Profiling in the Brain of a Transgenic Mouse Model of Alzheimer’s Disease Reveals Increased Prenylation of a Key Set of Proteins. ACS Chem Biol 17, 2863–2876 (2022). 10.1021/acschembio.2c00486

102 Cuddy, L. K. et al. Farnesyltransferase inhibitor LNK-754 attenuates axonal dystrophy and reduces amyloid pathology in mice. Mol Neurodegener 17, 54 (2022). 10.1186/s13024-022-00561-9

103 Hernandez, I. et al. A farnesyltransferase inhibitor activates lysosomes and reduces tau pathology in mice with tauopathy. Sci Transl Med 11 (2019). 10.1126/scitranslmed.aat3005

104 Cheng, S. et al. Farnesyltransferase haplodeficiency reduces neuropathology and rescues cognitive function in a mouse model of Alzheimer disease. J Biol Chem 288, 35952–35960 (2013). 10.1074/jbc.M113.503904

105 Jung, D. & Bachmann, H. S. Regulation of protein prenylation. Biomed Pharmacother 164, 114915 (2023). 10.1016/j.biopha.2023.114915

106 Liu, Z. et al. Membrane-associated farnesylated UCH-L1 promotes alpha-synuclein neurotoxicity and is a therapeutic target for Parkinson’s disease. Proc Natl Acad Sci U S A 106, 4635–4640 (2009). 10.1073/pnas.0806474106

107 Pan, J., Song, E., Cheng, C., Lee, M. H. & Yeung, S. C. Farnesyltransferase inhibitors-induced autophagy: alternative mechanisms? Autophagy 5, 129–131 (2009). 10.4161/auto.5.1.7329

108 Cuddy, L. K. et al. Stress-Induced Cellular Clearance Is Mediated by the SNARE Protein ykt6 and Disrupted by alpha-Synuclein. Neuron 104, 869–884 e811 (2019). 10.1016/j.neuron.2019.09.001

109 Roberts, P. J. et al. Rho Family GTPase modification and dependence on CAAX motif-signaled posttranslational modification. J Biol Chem 283, 25150–25163 (2008). 10.1074/jbc.M800882200

110 Prendergast, G. C. Actin’ up: RhoB in cancer and apoptosis. Nat Rev Cancer 1, 162–168 (2001). 10.1038/35101096

111 Schmidt, S. I., Blaabjerg, M., Freude, K. & Meyer, M. RhoA Signaling in Neurodegenerative Diseases. Cells 11 (2022). 10.3390/cells11091520

112 Tonges, L. et al. Rho kinase inhibition modulates microglia activation and improves survival in a model of amyotrophic lateral sclerosis. Glia 62, 217–232 (2014). 10.1002/glia.22601

113 Benarroch, E. What Is the Role of the Rho-ROCK Pathway in Neurologic Disorders? Neurology 101, 536–543 (2023). 10.1212/WNL.0000000000207779

114 Koch, J. C. et al. Safety, tolerability, and efficacy of fasudil in amyotrophic lateral sclerosis (ROCK-ALS): a phase 2, randomised, double-blind, placebo-controlled trial. Lancet Neurol 23, 1133–1146 (2024). 10.1016/S1474-4422(24)00373-9

115 Seijffers, R., Mills, C. D. & Woolf, C. J. ATF3 increases the intrinsic growth state of DRG neurons to enhance peripheral nerve regeneration. J Neurosci 27, 7911–7920 (2007). 10.1523/JNEUROSCI.5313-06.2007

116 Kim, Y. H. et al. A 3D human neural cell culture system for modeling Alzheimer’s disease. Nat Protoc 10, 985–1006 (2015). 10.1038/nprot.2015.065

117 Held, A. et al. iPSC motor neurons, but not other derived cell types, capture gene expression changes in postmortem sporadic ALS motor neurons. Cell Rep 42, 113046 (2023). 10.1016/j.celrep.2023.113046

118 Seczynska, M., Bloor, S., Cuesta, S. M. & Lehner, P. J. Genome surveillance by HUSH-mediated silencing of intronless mobile elements. Nature 601, 440–445 (2022). 10.1038/s41586-021-04228-1

119 Ge, S. X., Jung, D. & Yao, R. ShinyGO: a graphical gene-set enrichment tool for animals and plants. Bioinformatics 36, 2628–2629 (2020). 10.1093/bioinformatics/btz931

120 Hart, T., Komori, H. K., LaMere, S., Podshivalova, K. & Salomon, D. R. Finding the active genes in deep RNA-seq gene expression studies. BMC Genomics 14, 778 (2013). 10.1186/1471-2164-14-778

121 Stephens, M. False discovery rates: a new deal. Biostatistics 18, 275–294 (2017). 10.1093/biostatistics/kxw041

122 Roberts, B. et al. Fluorescent Gene Tagging of Transcriptionally Silent Genes in hiPSCs. Stem Cell Reports 12, 1145–1158 (2019). 10.1016/j.stemcr.2019.03.001

123 Roberts, B. et al. Systematic gene tagging using CRISPR/Cas9 in human stem cells to illuminate cell organization. Mol Biol Cell 28, 2854–2874 (2017). 10.1091/mbc.E17-03-0209

124 Labun, K. et al. CHOPCHOP v3: expanding the CRISPR web toolbox beyond genome editing. Nucleic Acids Res 47, W171–W174 (2019). 10.1093/nar/gkz365

125 Renthal, W. et al. Transcriptional Reprogramming of Distinct Peripheral Sensory Neuron Subtypes after Axonal Injury. Neuron 108, 128–144 e129 (2020). 10.1016/j.neuron.2020.07.026

